# Dynamic macromolecular composition and high exudation rates in *Prochlorococcus*

**DOI:** 10.1101/828897

**Authors:** Dalit Roth-Rosenberg, Dikla Aharonovich, Anne-Willem Omta, Michael J. Follows, Daniel Sher

**Author notes:** Materials and Correspondence Please send requests for materials or other correspondence to Daniel Sher.

## Abstract

Every living cell is composed of macromolecules such as proteins, DNA, RNA and pigments. The ratio between these macromolecular pools depends on the allocation of resources within the organism to different physiological requirements, and in turn affects biogeochemical cycles of elements such as carbon, nitrogen and phosphorus. Here, we present detailed measurements of the macromolecular composition of *Prochlorococcus* MIT9312, a representative strain of a globally abundant marine primary producer, as it grows and declines due to nitrogen starvation in laboratory batch cultures. As cells reached stationary stage and declined, protein per cell decreased by ∼30% whereas RNA per cell and pigments per cell decreased by ∼75%. The decline stage was associated with the appearance of chlorotic cells which had higher forward scatter (a proxy for cell size) but lower chlorophyll autofluorescence, as well as with changes in photosynthetic pigment composition. Specifically, during culture decline divinyl-chlorophyll-like pigments emerged, which were not observed during exponential growth. These divinyl-chlorophyll-like pigments were also observed in natural samples from the Eastern Mediterranean. Around >80% of the carbon fixed by *Prochlorococcus* MIT9312 (but not of a different strain, NATL2A) was released into the growth media as dissolved organic carbon under these laboratory conditions. Variations in RNA/protein indicate that, broadly defined, the macromolecular composition of *Prochlorococcus* MIT9312 is more similar to eukaryotic phytoplankton than to marine heterotrophic bacteria, possibly due to the significant investment in photosynthetic machinery of phototrophs.

## Introduction

Like every cell on Earth, phytoplankton and bacterioplankton are composed of several classes of biological macromolecules that function together to maintain the cells structure and activity (Geider and La Roche 2002, Finkel et al. 2016). Proteins are the most abundant macromolecule (in terms of mass), responsible for most of the metabolic functionality of the cells. DNA stores hereditary information, whereas RNA (as mRNA or other regulatory RNAs) accesses this stored data and (as rRNA and tRNAs) uses it to produce protein. Lipids form the membrane(s) that separate the cell from its environment and, in the case of photosynthetic organisms, host the molecular machinery used to harvest light for energy. In addition, the molecular machinery includes photosynthetic pigments and metabolites. Finally, various storage molecules (including some lipids, carbohydrates, polyphosphate and other storage molecules) allow cells to retain energy and elements for future use, and a host of other metabolites are involved in all forms of cellular function. The relative amount of each of these classes of macromolecules changes between different phytoplankton groups (Vargas et al. 1998, Geider and La Roche 2002, Finkel et al. 2016), and within a single type of organism in response to environmental conditions (Vargas et al. 1998, Liefer et al. 2019). For example, the relative amount of chlorophyll-a per cell changes in response to light intensity (e.g. (Moore et al. 1995)) and may also change in response to nutrient starvation (Lourenço et al. 1998, Liefer et al. 2019). This affects the ability of photosynthetic cells to harvest light and fix carbon, impacting cell physiology and likely the interactions of the cell with other organisms, e.g. through the release of fixed organic carbon (Dubinsky and Berman-Frank 2001). Similarly, it has been suggested that the number of ribosomes per cell is one of the factors determining cellular growth rate and is itself affected by temperature and nutrient (particularly phosphorus, P) availability (e.g. (Elser et al. 2003, Garcia et al. 2016, Martiny et al. 2016)). Thus, to some extent, the macromolecular composition of the cell is affected by, and may shed light on, the allocation of cellular resources to different functions. In addition, the basic building blocks of biological macromolecules such as amino acids and nucleic acids have distinct elemental composition, and the macromolecular and elemental composition of the cell are thus intimately linked (Geider and La Roche 2002, Finkel et al. 2016, Garcia et al. 2016, Liefer et al. 2019).

Here, we measure the cellular pools of major macromolecules (protein, DNA, RNA and pigments) in laboratory batch cultures of *Prochlorococcus*, a highly abundant pico-cyanobacterium which is responsible for ∼8.5% of the oceanic photosynthesis (Partensky and Garczarek 2010, Flombaum et al. 2013, Biller et al. 2014). *Prochlorococcus*, as a clade, is comprised of multiple genotypes, each different in its physiology, genome structure and oceanic niche (Johnson et al. 2006, Biller et al. 2014). Many aspects of the physiology of the *Prochlorococcus* clade have been intensively studied, ranging from its elemental composition (Bertilsson et al. 2003, Martiny et al. 2013) through its photo-physiology (Moore et al. 1995, Moore and Chisholm 1999, Ting et al. 2002, Steglich et al. 2003, Komatsu et al. 2016) to its system-wide transcriptomic response to changes in environmental conditions (e.g. (Martiny et al. 2006, Tolonen et al. 2006, Thompson et al. 2011, Aharonovich and Sher 2016). Yet, detailed analyses of the macromolecular composition of these cells are lacking. Strain MIT9312 represents the high-light adapted HL-II clade which is the most abundant clade in the surface waters of large parts of the ocean (Bouman et al. 2006, Johnson et al. 2006). We followed batch cultures of *Prochlorococcus* MIT9312 from exponential growth through to stationary stage, which occurred when the nitrogen (N) source in the media was depleted (Grossowicz et al. 2017). We chose to examine a nitrogen-limited batch culture across growth stages because, firstly, metaproteomic analyses show that N stress affects *Prochlorococcus* in large parts of the ocean (Saito et al. 2014). Secondly, significant information is available as to the physiological, transcriptomic and evolutionary responses of the cells to acute N starvation (e.g. (Steglich et al. 2001, Moore et al. 2002, Tolonen et al. 2006, Gilbert and Fagan 2011, McDonagh et al. 2012, Read et al. 2017, Domínguez-Martín et al. 2018, Berube et al. 2019, Szul et al. 2019)). Thirdly, long-term analyses of batch cultures may identify cellular physiological processes not seen when cells are growing exponentially or exposed to sudden or short-term nutrient starvation (e.g. (Christie-Oleza et al. 2017, Roth-Rosenberg et al. 2019)). Fourthly, nitrogen-stressed and starved cells have commonly been shown to accumulate or release DOC (as reviewed by (Thornton 2014)) with important implications for ecosystem interactions and the storage of potentially long-lived DOM in the ocean. (Hansell et al. 2009). Thus we characterized the changes in macromolecular composition, the specific complement of pigments and the rate of accumulation of organic carbon (both in particles and dissolved) from exponential growth through to stationary and decline phases.

In the following sections we describe the experimental methodology and the time-course of macro-molecular composition, pigments and TOC in the experiment. We discuss the changes in RNA:protein and its relationship to prior studies of eukaryotic phytoplankton and heterotrophic bacteria, the appearance of specific pigments in stationary phase which we have also observed in natural populations, as well as the accumulation of significant amounts of TOC over the course of the culture.

## Materials and Methods

### Growth conditions and experimental procedure

To follow the growth and macromolecular composition of *Prochlorococcus* over the various stages of laboratory batch culture, axenic *Prochlorococcus* MIT9312 cultures were grown in 1 liter bottles of Pro99 media (Moore et al. 2007) where the NH_4_ concentration was lowered to 100µM, leading to cessation of cell growth due to N starvation (Grossowicz et al. 2017). The experiment included 15 bottles, three of which were used for routine culture monitoring and the other 12 were used to collect samples at four time-points, corresponding to exponential growth (t=6 days), early stationary stage (t=10 days), late stationary stage (t=13 days) and culture decline (t=15 days). Samples were also collected from the starter culture to represent t=0. The number and timing of the samples collected for full analysis were selected in order to provide appropriate data for subsequent modeling (Omta et al. 2017). This experimental design was used to minimize the chance for contamination of the axenic cultures with heterotrophic bacteria. The cell numbers of the cultures used for macromolecular analysis were never statistically different from those of the cultures used for monitoring (Students t-test).

Prior to the beginning of the experiment, cells from a mid-exponential culture were counted by flow cytometry (FACSCantoII, BD), and diluted in the new media to an initial cell density of 10^6^/ml. Cultures were maintained under constant light (22 µE), at 24.5±1ºC. 1mM NaHCO_3_ was supplemented at three different time points (T_0_, T_7_, T_11_) to verify that the cultures are not carbon-limited (Moore et al. 2007, Grossowicz et al. 2017). Bulk culture fluorescence was measured in samples collected aseptically from the three monitoring bottles using a Carey Eclipse spectrofluorometer (Ex 440nm/Em 680nm). Samples for cell counting were fixed in 0.25% glutaraldehyde, kept in the dark for 10 minutes and transferred to a −80°C freezer for subsequent flow cytometry analysis.

### Measurements of macromolecular pools and photosynthetic pigments

At each of the five time-points selected, cells were collected by filtration. For protein measurements, cells were gently filtered on 0.22µm polycarbonate filters and kept in −80°C until extraction. For protein extraction, the filters were incubated in 500μl lysis buffer (50mM Tris pH6.8, 5mM EDTA, 2%SDS) on a lab rotator at 37°C for 20’. Following incubation, samples were sonicated in water bath for 10’, centrifuged at 12,000g in a table-top micro-centrifuge for 5’, and the supernatant liquid taken for measurement using the Bicinchoninic Acid assay (BCA, Sigma). Cells for measuring nucleic acids were collected similarly on 0.22 µm Supor-200 Membrane Disc Filters (25 mm; Pall Corporation) and preserved in storage buffer (40 mM EDTA, 50 mM Tris pH 8.3, 0.75 M sucrose) or RNA Save (Biological Industries) for DNA and RNA measurements, respectively. Samples were kept at −80°C until extraction. DNA was extracted from the filters according (Massana et al. 1997). Total RNA was extracted using the mirVana miRNA kit (Ambion, Austin, TX, USA) as described in with lysozyme treatment for better lysis (Tolonen et al. 2006), follow by removal of contaminating DNA from total RNA by Turbo DNase (Ambion). Nucleic acids quantity measured using Qubit 2.0 (Invitrogen). For pigment analysis, samples were collected on Glass Fiber filters (25 mm GF/F, Whatman) and kept in −80C until extraction. Glass fiber filters were used, rather than polycarbonate filters, as the latter are not compatible with many extraction methods for pigments. Nevertheless, preliminary experiments using a culture of *Prochlorococcus* MED4, which is similar in size to MIT9312, showed that with the vacuum pumps we used approximately 99.9% of the cells were retained on the GF/F membrane (Marmen et al. 2018), and thus the difference in pore size has minimal effect on the results. Cells were extracted in 100% methanol at 25°C for 2h. The cell extracts were clarified with Syringe filters (Acrodisc CR, 13 mm, 0.2 µm PTFE membrane, Pall Life Sciences), and kept at −20ºC until analysis by column chromatography on a C8 column (1.7 µm particle size, 2.1 mm internal diameter, 50 mm column length, ACQUITY UPLC BEH, 186002877) using an ACQUITY UPLC system equipped with a Photodiode Array detector (Waters). The chromatography protocol was based on LOV method (Hooker *et al.* 2005) modified for UPLC. A linear gradient was applied over 14 minutes, where solvent A was 70:30 100% methanol:0.5 M ammonium acetate, and solvent B was B. 100% methanol. The flow rate was 0.5 ml/min, and the column and injection heating were set to 50ºC and 30ºC, respectively. Photosynthetic pigments were identified based on their absorbance spectra and retention time, and were quantified using the following standards: Divinyl chlorophyll A (DVchlA), chlorophyll B (chlB), chlorophyll C (chlC), zeaxanthin (Zea), α-carotene (α-car), Divinyl Protochlorophyllide (MGDVP), Pheophorbide a, Pheophytin a and Chlorophyllide a. All standards were from DHI, Denmark.

### Measurements of inorganic nutrients and TOC

The GF/F filtrates (see above) were kept in polypropylene at −20ºC until analysis. For measurements, samples were adjusted to 25ºC and diluted 4-fold in DDW to reduce the salt level. NH_4_ (NH 3-N) was measured using an HI 96715 meter (Hanna instrument), and PO_4_ was measured by HI 96713 meter (Hanna instrument). In both cases, low concentration kits were used. For Total Organic Carbon (TOC) analysis, samples were collected into cleaned, pre combusted, acid washed (10% HCL) 40ml vials and HCl was added to remove dissolved inorganic C before preservation at −20ºC. Before analysis, samples were combusted at 680°C using catalytic (platinum) oxidation method in oxygen-rich environment. TOC analysis was performed using TOC-L analyzer (Shimadzu ASI-L Autosampler, Columbia, MD). For inorganic carbon (DIC), samples were collected in 14ml dark glass vials with screw caps and 75µl (0.05% v/v) saturated HgCl_2_ solution (Dickson et al. 2007) was added to the samples. For alkalinity, 60 ml samples were collected in glass vials. All samples were kept in 4°C until measurement. CO_2_ was extracted from the samples by acidification with phosphoric acid (H_3_PO_4_, 10%) using a custom, automated CO_2_ extractor and delivery system (AERICA by MARIANDA) using high grade N_2_ as a carrier gas, connected online with a LiCor 6252 IR CO_2_ analyzer. Measurements were calibrated using seawater Certified Reference Materials (CRMs) from Dickson’s lab. In order to check and correct for drift the CRM was run after every 4 samples.

### Model of *Prochlorococcus* resolving key macromolecular pools

We employed a mathematical model of the growth of a generalized axenic population of phytoplankton cells, which resolves the macromolecular pools measured in the experiments – protein, RNA, DNA and photosynthetic pigments, as well as N storage and dissolved organic carbon (Supplementary Figure S3). The model is formulated in terms of the flow and allocation of N in each macromolecular pool. In the model, proteins act as enzymes in the synthesis of RNA, DNA and Chl, whereas RNA acts as an enzyme for the synthesis of proteins. Key model parameters are constrained by optimizing the simulation of measured macromolecular composition. Carbon content of the modeled macromolecular pools is defined by their nitrogen content and known elemental ratios (Geider and La Roche 2002). Thus, the model predicts particulate organic carbon, so the observed total organic carbon in the media (TOC, the sum of cellular C and DOC) constrains the overall balance between photosynthesis and respiration. We fit the model to the experimental data presented below, using the Metropolis-Hastings algorithm (Metropolis et al. 1953). A more detailed description of the model, including equations, can be found in the supplementary information, and the model itself (written in FORTRAN) is available on https://github.com/AWO-code/DalitDaniel_Model.

## Results

### Dynamics of cell numbers and macromolecular composition during growth and nitrogen starvation

To determine to what extent the macromolecular composition of *Prochlorococcus* changes between the different physiological states of a batch culture, we grew strain MIT9312 in laboratory batch cultures where the N:P ratio of the media was set to 2, thus leading to cessation of growth due to N starvation (Grossowicz et al. 2017). The bulk culture fluorescence, often used to monitor phytoplankton growth in a non-invasive way, increased exponentially until day 10, after which the culture fluorescence declined rapidly, with no clearly observable stationary stage (Figure 1a). The decline of the culture fluorescence coincided with the reduction of soluble NH_4_ to below detection threshold (<10uM), while PO_4_ and DIC levels remained high throughout growth and decline (Figure 1c, d). Cell counts by flow cytometry also showed an increase in the cell numbers until day 10, with a growth rate (μ) of 0.353 day^-1^ (±0.0239) and doubling time of 1.968 days (±0.129). However, unlike the bulk culture fluorescence that declines rapidly after this day, the cell numbers remained relatively stable for an additional 3-4 days, before starting to decline. During this period, a distinct population of cells with low chlorophyll autofluorescence (low-fl) emerged, identified by flow cytometry (Figure 1b, 67% low-fl cells on day 13 and 88% on day 15, (Roth-Rosenberg et al. 2019). These low-fl (“chlorotic”) cells also stained much more weakly with Sybr Green which binds to nucleic acids (primarily dsDNA but also ssDNA and RNA, Supplementary Figure S1b). At the same time, an increase in Sybr-Green staining was observed in the high-fl cells, potentially related to the arrest of these cells in G2 stage (Supplementary Figure S1, (Zinser et al. 2009)). The median forward scatter, a proxy for cell size, increased for both the high-fl and low fl cells (Supplementary Figure S1c).

**Figure 1:**
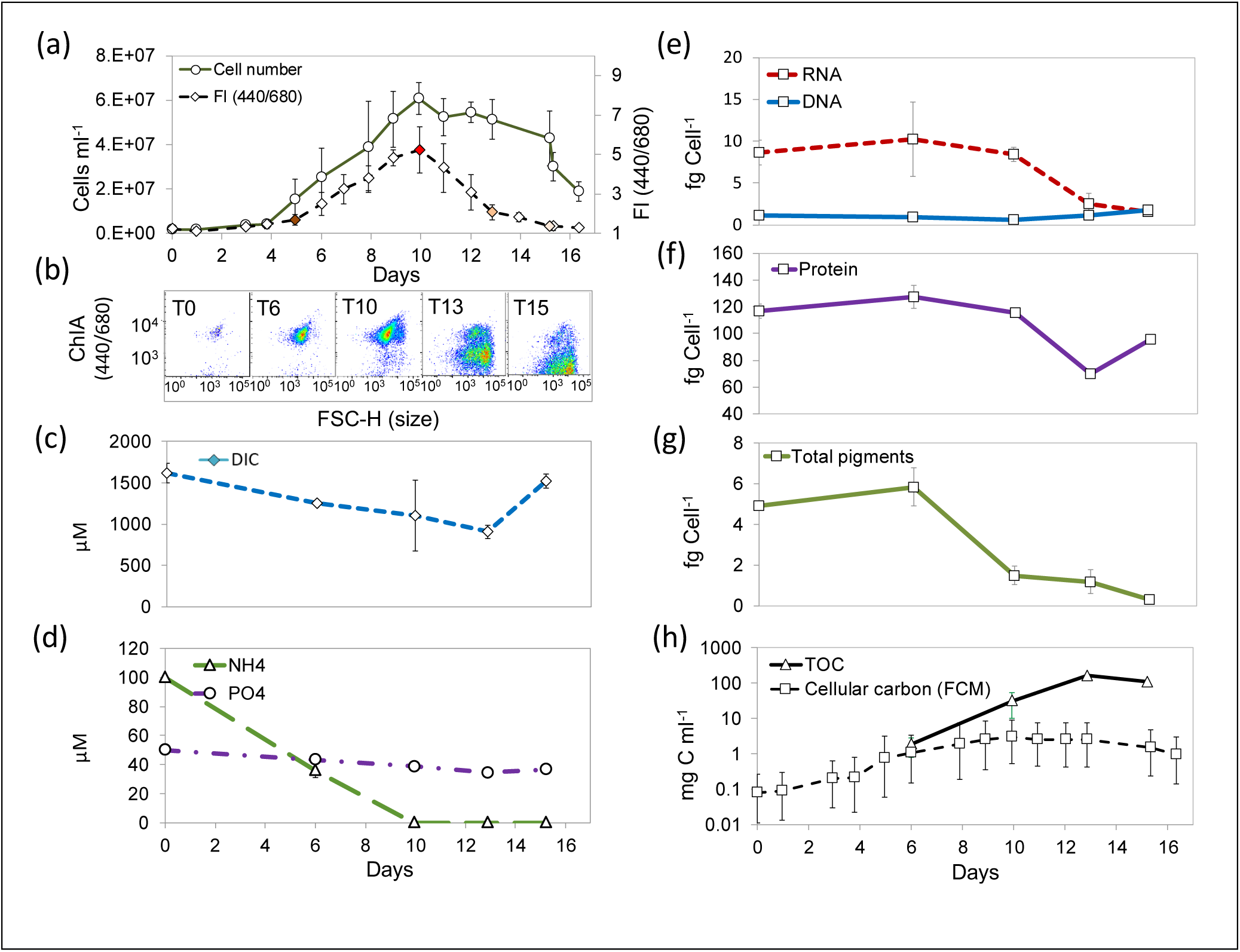
Dynamics of culture growth, plateau and death in batch *Prochlorococcus* MIT9312 cultures reaching stationary growth due to nitrogen starvation. (a) Cell numbers and fluorescence curves of MIT9312 population shows a log-phase growth until day 10 than stationary and decline of the culture. (b) Flow cytometry scatter gram of specific measurements points. (c-d) Analysis of external inorganic nutrients shows that NH_4_^+^ drops below detection limit, while PO_4_ and DIC are still available. Thus, the cultures enter stationary stage due to N starvation. Results are means and ranges of triplicate cultures. (e-g): Changes in the cell quotas of major macromolecules-DNA and RNA (e) proteins (f) and total pigments (g). While protein per cell is relatively stable, pigment and RNA pools drop as cells become starved and the culture declines. (h) Accumulation of large amounts of DOC. Triangles show measured Total Organic Carbon (TOC), squares show calculated C in cell biomass, based on flow cytometry counts and on per-cell C quotas. The error bars shown represents uncertainty due to the differences in estimates of per-cell C quotas between studies (Table 1), with the squares showing a value of 50 fg cell^-1^.

Based on the observed fluorescence curves (Figure 1a), we sampled cells during exponential stage (day 6), early stationary stage (day 10), late stationary stage (after bulk culture fluorescence had started to decline but while cell numbers were still stable, day 13) and culture decline (day 15) (Supplementary Table 1). An additional estimate of the macromolecular pools from exponentially-growing cells was obtained from the starter cultures and their properties provide the data for day 0. The per-cell content of proteins and RNA were relatively stable during exponential phase (0-10 days) but was reduced during the stationary and decline phases (days 13 and 15) (Figure 1e-f). The reduction in RNA per cell (a drop of approximately 75% between days 10 and 13) was much larger than that of protein per cell (10-30%). The total photosynthetic pigment concentration per cell started declining earlier than RNA or protein, with a drop of approximately 75% between days 6 and 10, as the cells moved from exponential growth to early stationary stage (Figure 1g).

### Changes in photosynthetic pigment composition across the different growth stages

The reduction in chlorophyll autofluorescence per cell suggests changes to the photosynthetic machinery of the cells as they enter stationary stage and during culture decline. Indeed, as shown in Figure 2a, in addition to a reduction in the total pigment quota per cell (Figure 1g), the ratio of some accessory pigments to divinyl chlorophyll A (DVchlA) changed over time. Divinyl chlorophyll B (chlB) and a chlorophyll-C like pigment (chlC) were reduced compared to DVchlA after day 10, whereas the ratio of zeaxanthin (and also α-carotene) to DVchlA increased after this day (Figure 2b). In addition, on day 10 and later, several UPLC peaks were observed close to that of DVchlA that differed from it in their retention time. These peaks were not due to overloading of the UPLC column (were not seen when larger amounts of pigments were injected) and were reproducibly observed in other cultures from multiple *Prochlorococcus* strains. One of these peaks (marked with an arrow in Figure 2c), which was especially prominent, had an absorption spectrum similar to that of DVchlA (Figure 2d) but was not one of the known degradation products (pheophytin-A eluted later, whereas phaeophorbide A, chlorophyllide A and divinyl protochlorophillide A all eluted much earlier, consistent with the loss of the phytol moiety). Assuming a similar molar absorption rate as DVchlA, this pigment (which may be a DVchlA’, an epimerization product of DVchlA, (Komatsu et al. 2016)) comprised as much as 21% of the total DVchlA pigments on day 15, at the end of the decline stage. Importantly, pigments with the same elution time and absorbance were repeatedly observed in samples collected from the oligotrophic Eastern Mediterranean Sea where *Prochlorococcus* contributes a significant fraction of the phototrophic biomass (Man-Aharonovich et al. 2010), as demonstrated in Supplementary Figure S2.

**Figure 2.**
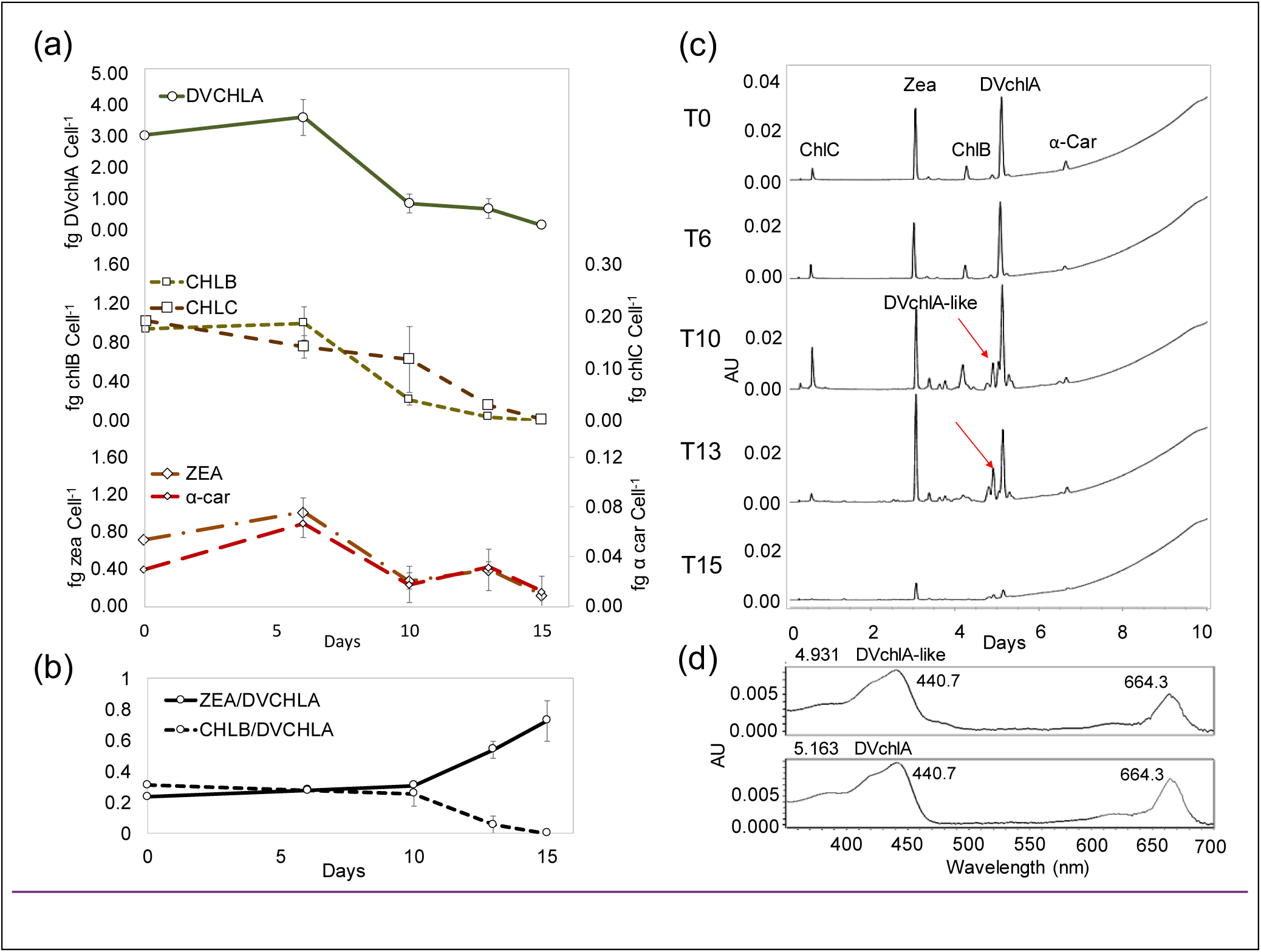
Pigments content and relative composition change during growth. (a) Changes in the major per-cell pigment quotas. Divinyl chlorophyll A - DVchlA, chlorophyll B - chlB, a chlorophyll C-like pigment - chlC, zeaxanthin - Zea, and α- carotene - α-car. (b) The relative abundance of zea to DVchlA increases, while that of chlB to DVchlA decreases. (c) UPLC chromatograms at different times show the appearance of additional pigments during stationary phase. The pigment marked with an arrow has an absorption spectrum similar to that of DVchlA (panel d), but differs from it in its retention time.

### Reproducible, high rates of DOC exudation in *Prochlorococcus* MIT9312

Part of the organic carbon fixed by *Prochlorococcu*s is released from the cell due to exudation and cell mortality. In natural settings this organic carbon provides sustenance for co-occurring heterotrophic bacteria (e.g. (Bertilsson et al. 2005, Agustí and Duarte 2013, Ribalet et al. 2015)). To assess how much organic C was released by *Prochlorococcus*, we measured the total organic C in our cultures, and compared it to the amount of C estimated to be in the particulate fraction based on the number of cells and the carbon quota of *Prochlorococcus* from previous studies (10-130 fg cell^-1^, Table 1). During exponential stage (day 6), the measured TOC was within the range expected based on cell numbers and C quotas. However, during the stationary and decline phases the amount of TOC was up to 23-fold higher than could be explained by cell biomass (assuming a maximal per-cell C biomass of 130 fg cell^-1^, Figure 1g, Table 1). This suggests a large amount of organic C was released into the growth media. To check if the high release of organic carbon is common to different *Prochlorococcus* strains, we repeated this experiment using both strain MIT9312 and strain NATL2A which belongs to the Low Light I clade (Biller et al. 2014). High extracellular concentrations of organic C (up to 13-fold higher than the predicted cellular biomass) were inferred again for MIT9312, whereas strain NATL2A that grew under the same N starvation conditions showed TOC values much lower than MIT9312, within the expected range of cell quota (Figure 3).

**Table 1:** *Prochlorococcus* Carbon biomass estimated in different studies, expanded from (Bertilsson et al. 2003)

| Strain | fg C cell<sup>-1</sup> | Analytical method | Reference |
| --- | --- | --- | --- |
| Natural population, Atlantic Ocean | 10-70 | X ray microanalysis | (Grob et al. 2013) |
| PCC9511, EQPAC, SB, GP2, SARG, NATL1 | 17-34 | X ray microanalysis | (Heldal et al. 2003) |
| PCC9511 | 17-38 | POC | (Claustre et al. 2002) |
| MED4 | 17-40 | POC | (Fu et al. 2007) |
| VOL7(MED4),<br>VOL8(MIT9515), VOL29,<br>VOL4 (MIT9312),<br>VOL1(MIT9215), UH18301 | 18-55 | POC | (Martiny et al. 2016) |
| MIT9301, MED4, NATL2A,<br>MIT9313 | 30-33 (HL),<br>45 (LLI),<br>80 (LLIV) | Macromechanical mass<br>sensor, assuming C is<br>50% of wet biomass | (Cermak et al. 2016) |
| MIT9301, MED4, MIT9313 | 44-51 (HL),<br>~180<br>(LLIV) | POC | (Becker et al. 2014) |
| MED4 | 46-61 | POC | (Bertilsson et al. 2003) |
| CCMP1378(MED4) | 49+/-9 | POC | (Cailliau et al. 1996) |
| Generic<br>(0.6 $\mu\text{m}$ diameter cell) | 53 | Cell size and volume-<br>carbon conversion<br>(470 fg C/$\mu\text{m}^3$) | (Campbell et al. 1994) |
| Natural population,<br>Atlantic Ocean | 56 | Cell size and volume-<br>carbon conversion<br>(325 fg C/mm<sup>3</sup>) | (DuRand et al. 2001) |
| Generic<br>(0.8 $\mu\text{m}$ diameter cell) | 59 | Cell size and volume-<br>carbon conversion<br>(220 fg C/mm<sup>3</sup>) | (Li et al. 1992) |
| MIT9211, MIT9215,<br>MIT9312, MED4, SS120,<br>MIT9302, MIT9303,<br>MIT9313 | 61-94 | Primary production,<br>potential<br>overestimation (does<br>not account for<br>respiration and<br>exudation) | (Moore 1997) |
| MIT9312 | 67-123 | Macromolecular pools | This study |
| MED4, MIT9401, SS120 | 78+/-27 | Dry weight, assuming C<br>is 50% of biomass | (Shaw et al. 2003) |
| Generic<br>(0.8 $\mu\text{m}$ diameter cell) | 124 | Cell size and volume-<br>carbon conversion<br>(294 fg C cell<sup>-1</sup>) | (Veldhuis and Kraay<br>1990) |

**Figure 3:**
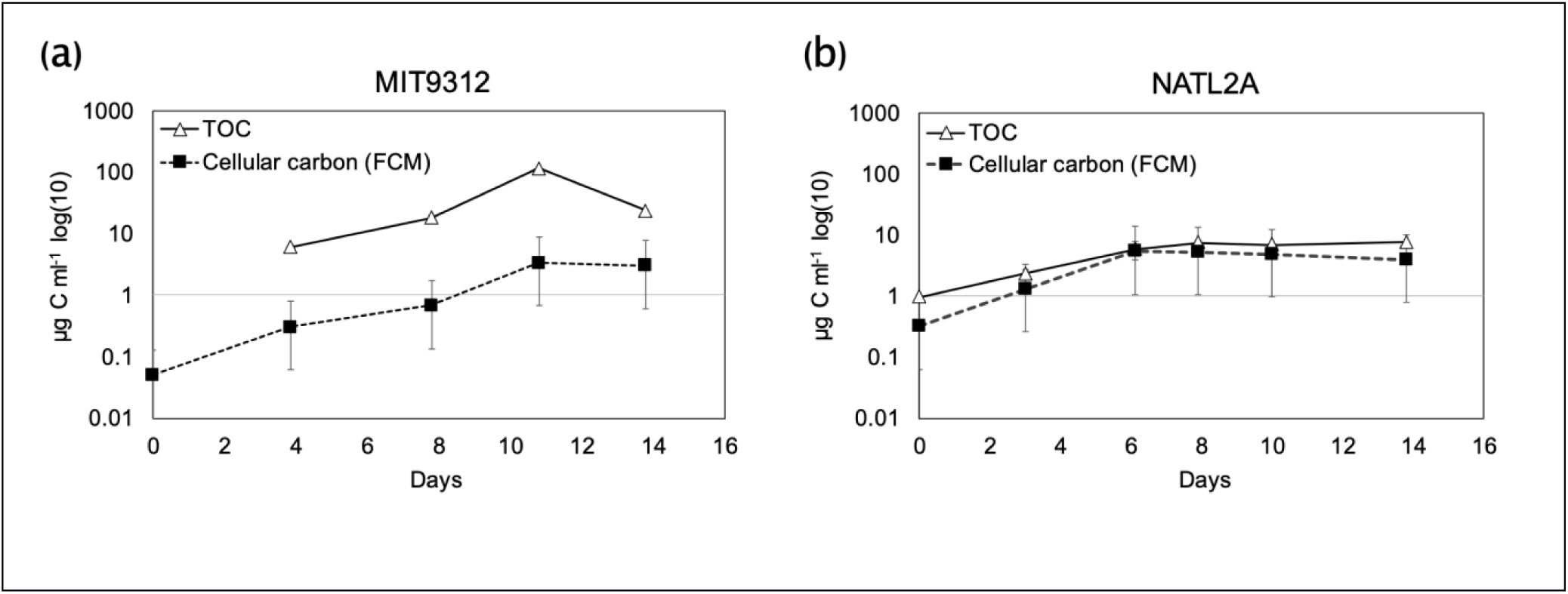
Differences in exudation rates between two axenic *Prochlorococcus* strains, MIT9312 (a) and NATL2A (b). The two strains were grown under identical conditions. Triangles show measured Total Organic Carbon (TOC), squares show calculated C in cell biomass, based on flow cytometry counts and on per-cell C quotas. The range shown represents uncertainty due to the differences in estimates of per-cell C quotas between studies (Table 1), with the square markers showing a value of 50 fg cell^-1^. Panel a) shows an independent experiment from that shown in Figure 1h, yet the high release of DOC is consistent.

## Discussion

In the following, we first detail some of the key assumptions and caveats, before going on to discuss three key results: the changing RNA:protein ratio, the appearance of signature pigments in stationary phase, and the very significant release of DOC into the medium under nitrogen starvation.

### Assumptions, Qualifications and Caveats

Quantitative measurements of cellular parameters, including macromolecular composition and elemental quotas, are fundamental to our understanding (and modeling) of microbial systems. However, measurements of the per-cell C quota of *Prochlorococcus*, performed using different techniques, range over an order of magnitude (Table 1, (Bertilsson et al. 2003)). In this study, we directly measured the cell quotas of DNA, RNA and photosynthetic pigments at different physiological stages of laboratory batch culture in *Prochlorococcus* MIT9312. With some assumptions, these measurements can be used to constrain upper and lower bounds of the total cell quotas of C, N and P (Supplementary text, Supplementary Table 1). Measuring these macromolecular pools requires less biomass and can be performed at higher throughput compared to measurements of total particulate or dissolved C and N (Supplementary text). Assuming a lower bound on the C:N ratio of 6 (Bertilsson et al. 2003, Martiny et al. 2013, Martiny et al. 2016), the cellular C quota we estimate (∼67-123 fg cell^-1^, Supplementary Table 1) is at the upper range of measurements from other studies (Table 1), and is driven primarily by the high (and mostly stable) measurements of protein per cell. These measurements are sensitive to the experimental design and the methodology used. For example, while the method we used for protein determination (BCA) is relatively insensitive to the effects of salts and detergents (if these are incorporated into the standard curve), the quantitative results can be affected by, for example, the protein standards used (e.g. Bovine Serum Albumin or Immunoglobulin). Similarly, our estimates of DNA per cell are about one-third those expected from cell number and the genome size of MIT9312 (1.7Mbp, (Kettler et al. 2007)). Experimental measurements of DNA per cell that are not consistent with genome size have been observed also in other studies (Zimmerman et al. 2014), and may be caused by differences in the DNA extraction efficiency across organisms or culture conditions. Finally, the *Prochlorococcus* population is not homogenous, containing at times two distinct sub-populations of cells, with different flow cytometry signatures (as discussed in detail below, see also Supplementary Figure S1 and (Coe et al. 2016, Roth-Rosenberg et al. 2019)). We did not measure the amount of the macromolecular pools in each sub-population separately, due to the difficulty in obtaining sufficient biomass from non-fixed cells sorted by fluorescence-activated cell sorting. With these uncertainties in mind, in the following sections we use the measurements of macromolecular pools at different growth stages, as well as the estimates of cellular C, N and P, to discuss the relationship between cell physiology, macromolecular pools and elemental budgets. Doing so, we note the need for an in-depth comparison of measurements between labs, growth stages and methods, perhaps along the lines of similar studies used to calibrate pigment measurements (e.g. (Claustre et al. 2004)).

### Dynamic changes in the macromolecular composition of *Prochlorococcus* in batch culture – changes in the average cell or sub-populations?

In our laboratory batch cultures, *Prochlorococcus* populations grew exponentially until they ran out of their nitrogen source (NH_4_), at which stage they stopped growing, entered a short stationary stage and finally died. During this time, the cultures lost their green color, the cell population became heterogeneous, and a sub-population of low-fl cells emerged, forming the majority of the population as the cultures decline (Figure 1a, Supplementary Figure S1a). The low-fl cells also stain weaker with the dye Sybr-green (Supplementary Figure S1), which binds both DNA and RNA (albeit with a higher affinity for DNA, (Martens-Habbena and Sass 2006)). This suggests that a significant part of the changes in the mean per-cell concentration of chlorophyll and RNA is caused by the increased fraction of these low-fl cells, rather than by a gradual change in the concentration of these pools per cell. The reduction in nucleic acids per cell is consistent with proteomic observations showing a 57% reduction in ribosomal proteins in a different strain of *Prochlorococcus*, SS120, under simulated nitrogen starvation (Domínguez-Martín et al. 2017). In the model cyanobacteria *Synechococcus elegantus* PCC7942 and *Synechocystis* PCC6803, the loss of culture chlorophyll (chlorosis) is part of a developmental program to generate resting stages, that can survive long-term starvation (e.g. (Sauer et al. 2001, Klotz et al. 2016)). In contrast, we have recently shown that *Prochlorococcus* cultures that become chlorotic cannot revive when nutrients are added, unless co-cultured with a heterotrophic bacterium (Roth-Rosenberg et al. 2019). Nevertheless, some of the chlorotic cells are still active (photosynthesize and take up NH_4_, (Roth-Rosenberg et al. 2019)). Thus, we interpret the changes observed in the macromolecular pools of the MIT9312 cultures, and primarily those that can be attributed to the low-fl (chlorotic) population, as physiological responses to N starvation that nevertheless are not sufficient to enable the cells to survive long-term nutrient stress.

In parallel to the observed loss of culture fluorescence and the appearance (and dominance) of low-fl cells, changes were also in the per-cell concentration and composition of the photosynthetic pigments (Figure 1h, Figure 2). Specifically, the mean relative amount of zeaxanthin increases compared to DVchlA, whereas the relative amount of chlB decreases (Figure 2b). Similar changes were observed in nutrient-replete batch cultures of other *Prochlorococcus* (strains MED4 and SS120) in response to increasing light intensities (Moore et al. 1995). One possible interpretation is that the changes in the pigment ratio as the cells enter stationary phase and start declining are due to increased light levels as many of the cells become chlorotic and the cultures become clear. An alternative interpretation is that the increase in relative abundance of zeaxanthin is a general stress response, as these pigments may act as antioxidants in cyanobacteria (Zhu et al. 2010). Indeed, an increase in the zeaxanthin-DVchlA ratio was observed upon nitrogen starvation also in an additional *Prochlorococcus* strain, SS120 (Steglich et al. 2001). The decrease in divinyl chlorophyll B can also be interpreted independently from a potential change in light levels. This reduction could simply be related to the deactivation of the photosystems, in agreement with previous studies that showed a decrease upon nitrogen starvation in photosynthetic activity (e.g. Fv/Fm, (Steglich et al. 2001, Lindell et al. 2002)), as well as in gene expression and protein amount of photosynthetic reaction center proteins (Tolonen et al. 2006, Domínguez-Martín et al. 2017). Alternatively, the reduction in DVchlB may be part of a cellular mechanism to reduce the use of N-containing pigments under N starvation, as chlorophylls contain nitrogen, whereas xanthophylls do not. This is consistent with other mechanisms whereby *Prochlorococcus* conserve N, including changes in the transcriptional profiles (Tolonen et al. 2006), use of shorter transcripts (Read et al. 2017) and, potentially, “thrifty” N use in proteins expressed in response to N starvation (Gilbert and Fagan 2011).

The emergence of the DVchlA-like pigment is currently unexplained, although this pigment may be an epimerization product of DVchlA (DVchlA’). DVchlA has been previously observed in several *Prochlororoccus* strains (Komatsu et al. 2016), and in our study the DVchlA-like pigment is associated with culture decline. Generally speaking, harvesting cells from late exponential stage or early stationary stage is a common practice when high biomass is required, yet as discussed above the physiology of the cells can change during stationary phase in comparison with exponentially-growing cells. Additionally, it is currently unclear to what extent processes observed in late stages of laboratory batch cultures, such as nutrient starvation and the subsequent changes in cell physiology, occur in nature. Nevertheless, the observation that a pigment with the same retention time and absorption spectrum as the DVchlA-like pigment observed at-sea suggests that at least some of the cells in nature may be undergoing processes similar to those we observe in the N-starved laboratory batch cultures.

The high-fl cells also exhibit a slight increase in mean forward scatter, a proxy for cell size (Supplementary Figure S1). This occurs despite the overall reduction in RNA, pigments and (to a lesser extent) protein (Figure 1e, f), which are the major N-containing macromolecular pools. Assuming the increase in forward scatter represents an increase of approximately 1.2-1.5-fold in cell biomass (Roth-Rosenberg et al. 2019), this suggests that the cells are accumulating primarily C-rich macromolecules such as storage carbohydrates and lipids, leading to an increased C:N ratio. A similar response to nitrogen starvation has been predicted by mathematical models of N-starved cells (Grossowicz et al. 2017) and observed in *Prochlorococcus* (Lindell et al. 2002) as well as in other phytoplankton species (Breuer et al. 2012, Liefer et al. 2019). If these processes occur in nature, they could impact the C:N ratio of DOM released from dead *Prochlorococcus* cells (e.g. (Agusti and Sanchez 2002, Llabrés et al. 2011, Ribalet et al. 2015)) as well as POM exported by these cells (Richardson and Jackson 2007, Lomas and Moran 2012, Zhao et al. 2017).

### The macromolecular composition of *Prochlorococcus* MIT9312 is more akin to eukaryotic phytoplankton than to heterotrophic bacteria

Phytoplankton and heterotrophic bacteria are inherently different in their life histories, and this likely determines and constrains the relative allocation of energy and elements to different molecular functions. Phytoplankton need to allocate resources to the photosynthetic and carbon fixation machineries and are likely less constrained by the availability of carbon (which can be derived from photosynthesis) compared to other elements such as nitrogen. Eukaryotic phytoplankton are also often larger than bacteria, and thus are less constrained by the internal volume of the cell. Heterotrophic bacteria, in contrast, do not need to allocate resources to photosynthesis but are potentially limited by C and by cell size. As shown in Figure 4, in combination with measurements of macromolecular pools in marine microorganisms available from the literature, our data supports the idea phytoplankton and bacteria allocate resources differently.

**Figure 4.**
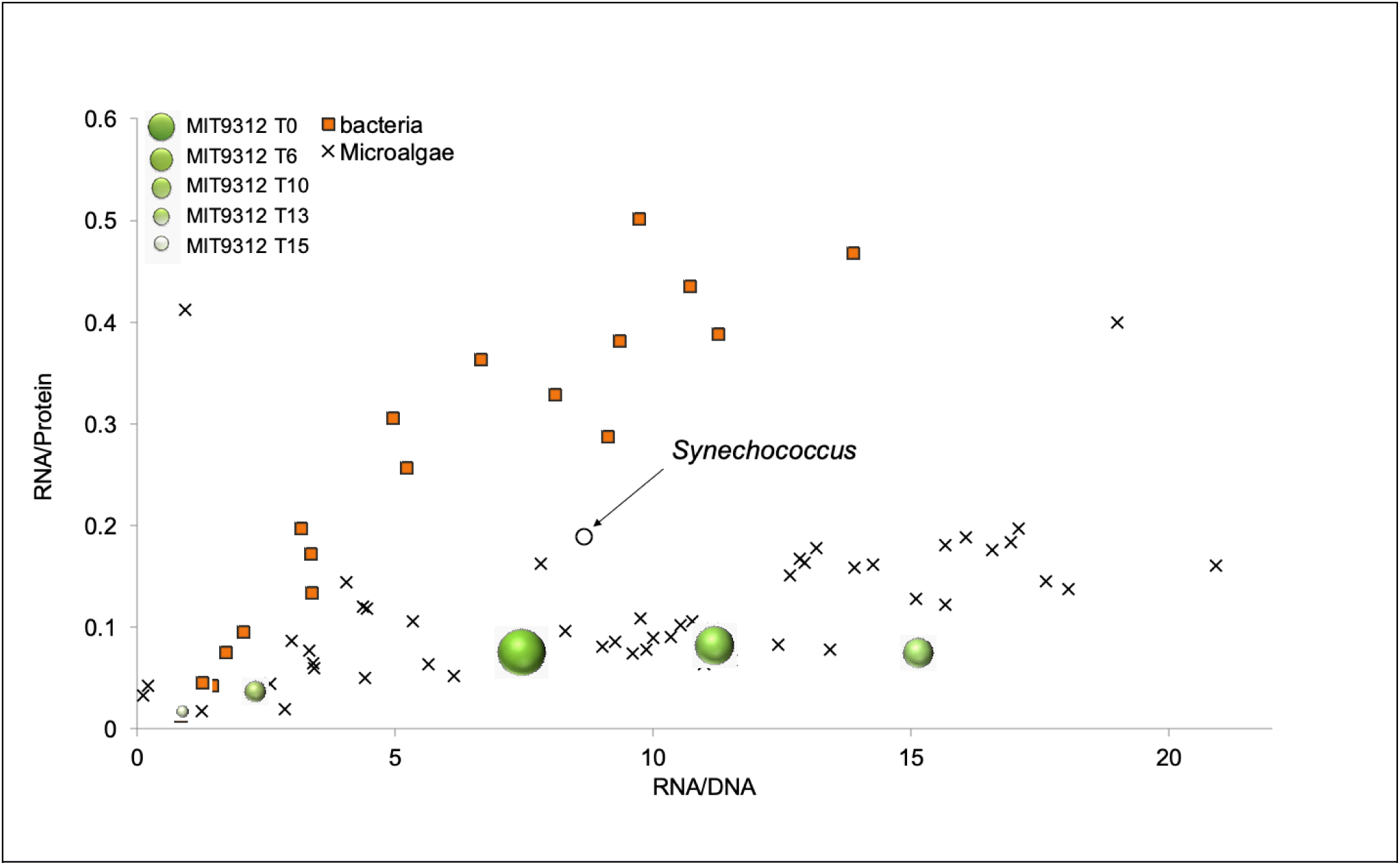
RNA/DNA and RNA/protein compared to those of other phytoplankton and bacteria. The ratios in phytoplankton (from (Finkel et al. 2016)), presented in cross markers, show a high diversity of RNA/DNA, while relatively low diversity of RNA/Proteins compositions. Heterotrophs bacteria however (squares) are highly diverse in both RNA/Proteins and RNA/DNA composition (Churchward et al. 1982, Neidhardt et al. 1990, Zimmerman et al. 2014). *Prochlorococcus* MIT9312 at different growth phases (spheres of different sizes) are in fact similar to microalgae rather to bacterial macromolecule composition. The most phylogenetically related cyanobacteria, *Synechococcus* WH7803, marked in circle, shows similarity to MIT9312 in the RNA/DNA but not in the RNA/Protein ratios.

Figure 4 shows that both heterotrophs and phototrophs exhibit a linear relationship between RNA and protein content. The relationships are consistent across taxonomic variations but consistently differ between phototrophs and heterotrophs. Generally speaking, phytoplankton have a lower RNA/protein ratio than heterotrophic bacteria with the same RNA/DNA ratio. Assuming that the ribosomes of phytoplankton and bacteria have similar maximal rates of protein production (chain elongation), this suggests that phytoplankton ribosomes are working at a higher relative capacity. In phytoplankton, a significant amount of protein production (up to ∼50% of the proteome, (Zavřel et al. 2019)) needs to be invested in maintaining the photosynthetic apparatus, for example in the replacement of the core proteins. An alternative (but non-exclusive) explanation is that heterotrophic bacteria (in particular copiotrophs) maintain more ribosomes than needed, using the “spare” production capacity to allow rapid response to changes in growth conditions. Such a strategy has been demonstrated experimentally for *E. coli* (e.g. (Li et al. 2018)). Notably, some bacteria, such as *Psychrobacter Mor119, Halomonas Hal146, Vibrio Vib2d* and *Ruegeria Oce241*, are more similar in the RNA/protein ration to phytoplankton (Figure 4). The reason for this similarity, which may represent a lower ability to allocate ribosome resources to rapid changes in gene expression, is unclear, and may depend on the specific experimental conditions employed in their study (Zimmerman et al. 2014). While *Prochlorococcus* is a bacterium (prokaryote), its RNA/protein and RNA/DNA ratios at different stages of the growth curve all fall within the range of those from eukaryotic phytoplankton. The single data point from *Synechococcus* is also closer to other phytoplankton (Figure 4). This could be the result of both the need to invest in maintaining the photosystem and, potentially, that *Prochlorococcus* may have a lower ability compared to fast-growing heterotrophs to “ramp up” their translation in order to rapidly respond to changes in environmental conditions.

Nevertheless, *Prochlorococcus* may still maintain some unused ribosome capacity. In our study, protein per cell remained relatively stable, with a decline of no more than ∼30% as the culture declined. RNA per cell, in contrast, dropped by more than 80% over the same period (Figure 1). The decline in RNA per cell also started earlier. This suggests either that processes resulting in the loss of protein (e.g. degradation, exudation or excretion) were strongly reduced in stationary and decline phase cells, or that the actual rate of protein production per ribosome increased in declining cultures compared to exponentially-growing ones. These two explanations - decrease in loss processes or increase in ribosome efficiency - are not mutually exclusive, but evidence is lacking for either of them in most organisms, including *Prochlorococcus*.

### DOC accumulation and its potential mechanism

One of the surprising observations of this study was the high rate of accumulation of DOC during the exponential growth of MIT9312. By the time the cultures reached stationary stage (day 13 in Figure 1h and day 11 in Figure 3a) the particulate organic carbon (i.e. cell biomass) was only 3-7% of the total organic carbon (considering the two experiments shown in Figure 1h and 3h, and assuming a cell C quota of 120 fg cell^-1^). This suggests that 80-85% of the organic carbon fixed by *Prochlorococcus* in two separate experiments was released as dissolved organic carbon. Previous experimental studies where DOC release was measured using ^14^C as a tracer suggested that 2-24% of the primary productivity is released as DOC (Bertilsson et al. 2005, Lopez-Sandoval et al. 2013), although higher loss values have also been reported (up to ∼60%, (Szul et al. 2019)). Indirect assessments suggest that a somewhat higher fraction of the fixed carbon is lost from the cells, respired as CO_2_ or released as DOC. Firstly, a comparison of carbon uptake through photosynthesis and growth rate in *Prochlorococcus* MED4 over a diel cycle suggested that ∼30% of the fixed carbon is released or respired (Zinser et al. 2009). Secondly, the expected growth rate of MIT9312 based on the calculated yield of photosynthesis (taking into account published values for illumination, cross section, efficiency and chlorophyll/ <u>per</u> cell, (Moore and Chisholm 1999)) is almost two-fold higher than measured (expected μ=0.64 day^-1^, compared to 0.35±0.02 day^-1^ measured during exponential stage, see Supplementary Information for more details). Both of these observations are consistent across phytoplankton species representing the difference between gross and net primary productivity. The differences between our results, based on accumulation of DOC over time, and estimates of exudation based on ^14^C partitioning between the particulate and dissolved phases, may also be due to loss of volatile substances during the acidification and subsequent venting of inorganic ^14^C during radioactive incorporation experiments. Indeed, *Prochloroccoccus* produce several volatile substances (e.g. organic acids, organohalogens, methane and isoprene, (Shaw et al. 2003, Bertilsson et al. 2005, Hughes et al. 2011, Bižić et al. 2020)), although to what extent these substances would be lost from the media during the protocol for measuring ^14^C incorporation is unclear.

To perform a careful and comprehensive budget, we used a mathematical model of a generalized phytoplankton to test whether the exudation rates we observed are feasible, given what we know about the underlying organismal physiology. The model represents the carbon and nitrogen budgets of, and flow between, the macromolecular pools that were measured (Supplementary Figure S3). The experimental measurements of these macromolecular pools were then used to directly fit the model to the data and constrain unknown parameters. The model is able to reproduce the change in cell density, the uptake of inorganic N and the reduction in RNA and chlA after the stationary stage (Supplementary Figure S4). The model captures the decline in protein in late stages of the culture, though the timing was earlier than observed. Importantly, the model requires a significant accumulation of DOC for mass balance, consistent with the large observed increase in TOC (Supplementary Figure S3-S5, Supplementary Table S2). The constrained model values of C fixation per unit chlA required to allow the model to produce the observed accumulation of TOC were reasonable; typically within the range of the maximum photosynthesis measured for *Prochlorococcus* (up to 10-20mol C [g DVchlA]^-1^day^-1^, (Moore and Chisholm 1999, Bruyant et al. 2005, Felcmanová et al. 2017)).

While the model was able to fit the data with reasonable parameter values, in order to simulate the high observed accumulation of TOC in the decline phase of the culture, the model had to either significantly increase the per chlorophyll carbon fixation rate *or* significantly reduce the basal metabolic respiration rate during the decline phase (Supplementary Figure S5). Such an increase in photosynthetic efficiency in cells entering chlorosis seems counter-intuitive. It might potentially be a response to a reduction in shelf-shading, (Felcmanová et al. 2017), but such a phenomenon has not, to the best of our knowledge, been described previously. A decrease in respiration rates in relation to growth rate as the cells enter chlorosis (Fang et al. 2019) is perhaps a more plausible hypothesis. The current model cannot distinguish between these two possibilities without further experimental constraints.

The DOM released by *Prochlorococcus* can include, in addition to C, also other elements such as N and P. We did not measure the elemental stoichiometry of the DOM but the difference between the N concentration in the initial media and the inferred N content of cell biomass under N-starved conditions represents organic N lost from cells that is refractory to *Prochlorococcus*. At day 10, peak growth in our experiment, the inferred cellular N accounted for 79% of the total N in the media. At this point, NH_4_ in the media was completely depleted suggesting that ∼20% of the N had already been released from the cells in a form that is not available for re-utilization. Thus, while C rich, the DOM released by *Prochlorococcus* MIT9312 contains significant amounts of other elements, likely in the form of proteins, amino acids, DNA, RNA and nucleotides. This is consistent with studies in *Synechococcus* (Christie-Oleza et al. 2015, Christie-Oleza et al. 2017, Zhao et al. 2017) and with the observed increase in amino acid metabolism following incubation of natural microbial communities with *Prochlorococcus*-derived DOM (Sharma et al. 2014). A similar calculation indicates that only ∼3.2% of the total P in the media was found in cellular biomass, while the PO_4_ concentration in the media dropped from 50μM to 34-36μM, a reduction of ∼30%. This could reflect unmeasured pools of intracellular pools of phosphorus (in addition to RNA, DNA and phospholipids), or that a significant amount of P has been converted from PO_4_ to extracellular DOP, or a combination of the two. Previous work (Rhee 1973, Liefer et al. 2019) suggests that the former is likely an important component of this un-accounted for P.

Several studies have suggested that high rates of DOC release occur in oligotrophic regions of the oceans, and that DOC, rather than POC, contributes significantly to carbon export from the photic zone in these areas ((Guyennon et al. 2015, Roshan and DeVries 2017), but see also (Iuculano et al. 2017)). It is tempting to speculate that *Prochlorococcus* may be at least partially responsible for this DOC accumulation, based on the high rates of DOC production observed here by *Prochlorococcus* MIT9312, a member of the HL-II clade that numerically dominates large areas of the oligotrophic ocean (Bouman et al. 2006, Johnson et al. 2006). In support of this speculation, DOM released by *Prochlorococcus* in the surface waters of the Pacific Ocean (which was dominated by the HL-II clade) has been suggested to provide as much as 75% of the daily photosynthetic organic C production (Ribalet et al. 2015), although it is unclear to what extent mortality compared to exudation feeds this organic pulse. It has also been suggested that DOM derived from *Prochlorococcus* contributes a significant amount to deep sea DOM (Zhao et al. 2017). We note, however, that DOC production rates may differ based on the physiological states of the cells and the surrounding nutrient conditions. For example, similar to our study, Bertilsson and co-workers also noted differences between two strains of *Prochlorococcus*, MIT9312 and MED4, with the former releasing approximately twice as much DOC (Bertilsson et al. 2005), and Becker and co-workers documented differences in the composition of released DOM (Becker et al. 2014). This could also depend on the mechanism whereby the DOM is released from the cells, namely passive leakage from the cells (exudation), active release (excretion) or lysis of dead cells (reviewed by (Thornton 2014)). Several studies have indicated that up to ∼20% of the cells in exponentially-growing *Prochlorococcus* cultures may be dead (Agusti and Sanchez 2002, Hughes et al. 2011). Future work is required to determine to what extent DOM production changes between strains and across culture conditions, what are the mechanisms of this process, and to what extent assessments of DOM release under lab conditions can be extended to the ocean.

## Conclusions

As laboratory batch cultures of *Prochlorococcus* MIT9312 grow exponentially, become N-starved and decline, the macromolecular composition of the cell changes. Cell quotas of RNA and photosynthetic pigments decline faster than proteins, and the cells likely accumulate C-rich storage molecules. At the same time, we observe a high production of extracellular DOC (DOM), primarily at the late stages of culture, which we propose is partly a result of a metabolic shift in the cells. This could either be an increased photosynthetic rate and/or a decreased respiration, although we cannot distinguish between these two possibilities here. These observations reflect the population mean, yet the dynamics may be different between different cellular sub-populations which are implied by flow cytometry.

While these results were observed in laboratory batch cultures, the presence of pigments in samples from the Eastern Mediterranean that are similar to those associated with culture decline in the lab, suggest that at least some of the cells in the oceans are undergoing similar changes in macromolecular structure, possibly due to N starvation, and are potentially releasing significant amounts of DOC. More generally, we propose that viewing cells through the “lens” of their macromolecular composition can help shed light on basic aspects of resource allocation, highlighting, for example, that prokaryotic phototrophs such as *Prochlorococcus* are more alike in this respect to eukaryotic phototrophs than to the heterotrophic bacteria to which they are more closely related evolutionarily.

## Acknowledgements

We thank Hila Elifantz, Ilana Berman-Frank, Eyal Geisler and Edo Bar-Zeev for the TOC analyses and Jack Silberman for the DIC analysis. This study was supported by the Gordon & Betty Moore Foundation (grant number GBMF #3778, to MJF), by the Human Frontiers Science Program (grant number grant RGP0020/2016, to DS) and by the United States-Israel Binational Science Foundation (Grant number 2010183 to DS and MJF and grant number 2016532 to DS).

## Author contributions

All authors conceived the study and designed the experiments. DR and DA performed the experiments and analyzed the samples. AWO performed the modeling analysis. DR and DS wrote the manuscript with contributions from all authors.

