## Supplementary Information for "Dynamic macromolecular composition and high exudation rates in *Prochlorococcus*"

### Constraining *Prochlorococcus* C and N quotas from measurements of macromolecular pools

Measuring the elemental composition of *Prochlorococcus* cells is important in order to understand their impact on biogeochemical cycles, yet performing such measurements is non-trivial. Some measurement techniques, such as CHN analysis, require significant biomass, which may not be easy to collect. For example, in order to obtain 5 milligrams of *Prochlorococcus* cells (the minimum requested by many analysis units), and assuming a dry weight of 100 fg cell$^{-1}$ (see Table 1 and the main text), $5\times10^{10}$ cells are needed. This corresponds to ~50 liters of an early exponential phase culture under our experimental setup, where cells are inoculated at ~$10^6$ cells/ml (e.g. Figure 1, (Sher et al. 2011, Aharonovich and Sher 2016), see also (Martiny et al. 2016)). While low compared to some other experiments, these cell initial densities are still 2-10 fold higher than conditions observed in nature even at times where *Prochlorococcus* are numerically dominant (e.g. (Malmstrom et al. 2010, Roth-Rosenberg et al. 2019), see also (Christie-Oleza et al. 2017)). For this reason, elemental composition is often measured in late exponential or early stationary stage cultures, potentially missing the dynamics described on this study (e.g. (Cailliau et al. 1996, Claustre et al. 2002, Bertilsson et al. 2003, Fu et al. 2007, Becker et al. 2014, Martiny et al. 2016). Other methods, e.g. based on X-ray analysis (Heldal et al. 2003, Grob et al. 2013) or micromechanical mass sensors (Cermak et al. 2016), are not widely available or amenable to high-throughput. Finally, estimations of cell biomass from flow cytometry require a conversion factor, which changes between studies (e.g. (Veldhuis and Kraay 1990, Li et al. 1992, Campbell et al. 1994, DuRand et al. 2001)). As a result, there is an order-of-magnitude range in the estimates of per-cell C biomass in *Prochlorococcus* (Table 1).

In contrast to these methods, measuring protein, DNA and RNA concentrations is relatively straightforward, requires instrumentation common to many biology labs, and can be adapted to low biomass and high throughput (e.g. by using a microplate reader). Like any analytical method, these also suffer from drawbacks and potential biases, as described in the main text. Nevertheless, with several assumptions, such measurements can be used to estimate the elemental quotas of the cells. Supplementary Table S1, below, shows the measured macromolecular pools at the different growth stages (in fg cell$^{-1}$), and the calculated masses of C and N for each pool using the known elemental compositions of each pool (Geider and La Roche 2002). To the summed C pools we add a value of 11.79 fg cell$^{-1}$ in membranes, based on the calculations presented in the supplementary information of Waldbauer et al (Waldbauer et al. 2012). In the paragraphs below, we discuss inferences that can be tentatively made from these measurements about the C, N and P budgets of the cells at different stages of the batch culture, comparing the estimates of elemental quotas to direct measurements using different methodologies from previous studies.

Assuming all of the N is found in the measured pools (protein, Chlorophyll, DNA and RNA), we obtain N quotas of 11.8-22.3 fg cell$^{-1}$. The values during late stationary and decline stages are within the range based on one study using X-ray microanalysis (1.5-18 fg cell$^{-1}$, (Grob et al. 2013)), but are higher than other studies using CHN analysis (3-9 fg cell$^{-1}$ in (Martiny et al. 2016), 3-8 fg cell$^{-1}$ in (Fu et al. 2007)). In contrast, the estimates we obtain during early exponential growth at low cell densities are somewhat higher than these values, perhaps reflecting differences in physiology related to cell density. We note that one study using X-ray microanalysis reported lower values (2.9-4.5 fg cell$^{-1}$, (Heldal et al. 2003)), but it also reported relatively low C quotas per cell as well as a relatively high C:N ratio of ~8.5-9.5.

While the N:P and C:P ratios of *Prochlorococcus* are highly variable, the C:N ratio is typically 6-7 (e.g. (Bertilsson et al. 2003, Martiny et al. 2013, Martiny et al. 2016)). Thus, assuming all of the N per cell has been accounted for as described above, a lower boundary for the cellular C quota can be calculated by multiplying the N quota by 6, resulting in a C quota of ~71-134 fg $cell^{-1}$. This value is high but within the range of previous estimates (Table 1). Comparing the calculated C quota with the sum of the measured pools yields a conservative estimate of the contribution of storage pools such as lipids and carbohydrates (e.g. glycogen), ranging from ~17-39 fg $cell^{-1}$ or 24-31% of the cell C (compare with, e.g. (Liefer et al. 2019)). This is a conservative estimate because the C:N ratio is likely to increase, especially in late stationary and declining cells where we observe an increase in the forward scatter of the cells (Supplementary Figure S1), potentially reflecting an increase in cell size driven by the accumulation of C storage molecules.

Finally, summing the P quotas in DNA and RNA, and adding 0.06 fg $cell^{-1}$ P from phospholipids (Waldbauer et al. 2012), yields and calculated P quota of 0.38-1.07 P fg $cell^{-1}$. This is within the range of analyses using measurements of particulate organic P (0.4-0.9 fg $cell^{-1}$ according to (Fu et al. 2007), 0.2-1.8 fg $cell^{-1}$ according to (Martiny et al. 2016)), and somewhat higher than estimates based on X-ray microanalysis (0.03-0.4 fg $cell^{-1}$ according to (Grob et al. 2013) and 0.25-0.49 fg $cell^{-1}$ according to (Heldal et al. 2003)).

Based on these calculated P quotas, the estimated N:P ratio of the cells is between ~20-22 during the exponential and early stationary stages, increasing to 30-43 during the late exponential and decline stages. These values are within the ranges described previously in batch cultures during exponential growth (Martiny et al. 2016) and P-starved cultures (Bertilsson et al. 2003), as well as for field populations from the oligotrophic oceans (Grob et al. 2013, Martiny et al. 2013). However, an increase in the N:P ratio as the cells transition into stationary stage is unlikely, given that stationary stage was due to N starvation. It is more likely that, given the high level of flexibility in the C:P and N:P ratios of *Prochlorococcus* cells (Bertilsson et al. 2003, Martiny et al. 2013, Martiny et al. 2016), our estimates of the P quotas are lower than the actual values. The additional P in the cells might be in the form of polyphosphate bodies or extracellular P reserves (Zubkov et al. 2015). Future experiments where these measurements are coupled with direct estimates of total P are required for this.

**Supplementary Table S1** – measured macromolecular pools and estimated elemental composition of *Prochlorococcus* MIT9312 cells during batch culture.

*Table S1*. **C and N estimation**

| | C and N content (fg cell$^{-1}$)[a] | | | | | | | | | | | | | | |
|---|---|---|---|---|---|---|---|---|---|---|---|---|---|---|---|
| **Days** | **0** | | | **6** | | | **10** | | | **13** | | | **15** | | |
| **Protein** | 116.88 | ± | 0.00 | 127.39 | ± | 19.74 | 115.55 | ± | 23.22 | 70.14 | ± | 24.93 | 95.81 | ± | 11.72 |
| $C_{Pro}$ | 59.61 | ± | 0.00 | 64.97 | ± | 10.07 | 58.93 | ± | 11.84 | 35.77 | ± | 12.72 | 48.86 | ± | 5.97 |
| $N_{Pro}$ | 18.70 | ± | 0.00 | 20.38 | ± | 3.16 | 18.49 | ± | 3.71 | 11.22 | ± | 3.99 | 15.33 | ± | 1.87 |
| **RNA** | 8.64 | ± | 4.03 | 10.23 | ± | 5.33 | 8.46 | ± | 2.75 | 2.46 | ± | 1.37 | 1.51 | ± | 0.58 |
| $C_{RNA}$ | 2.94 | ± | 1.37 | 3.48 | ± | 1.81 | 2.88 | ± | 0.94 | 0.84 | ± | 0.47 | 0.51 | ± | 0.20 |
| $N_{RNA}$ | 1.34 | ± | 0.62 | 1.59 | ± | 0.83 | 1.31 | ± | 0.43 | 0.38 | ± | 0.21 | 0.23 | ± | 0.09 |
| **DNA** | 1.16 | ± | 0.15 | 0.91 | ± | 0.21 | 0.56 | ± | 0.32 | 1.09 | ± | 0.14 | 1.80 | ± | 0.55 |
| $C_{DNA}$ | 0.42 | ± | 0.05 | 0.33 | ± | 0.08 | 0.20 | ± | 0.11 | 0.39 | ± | 0.05 | 0.65 | ± | 0.20 |
| $N_{DNA}$ | 0.19 | ± | 0.02 | 0.15 | ± | 0.03 | 0.09 | ± | 0.05 | 0.17 | ± | 0.02 | 0.29 | ± | 0.09 |
| **DVchlA** | 3.21 | ± | 0.00 | 3.89 | ± | 0.64 | 1.24 | ± | 0.75 | 1.53 | ± | 0.50 | 0.27 | ± | 0.18 |
| $C_{DVchlA}$ | 2.38 | ± | 0.00 | 2.88 | ± | 0.47 | 0.92 | ± | 0.55 | 1.13 | ± | 0.37 | 0.20 | ± | 0.13 |
| $N_{DVchlA}$ | 0.20 | ± | 0.00 | 0.25 | ± | 0.04 | 0.08 | ± | 0.05 | 0.10 | ± | 0.03 | 0.02 | ± | 0.01 |
| $C_{lipid}$[b] | 11.78 | | | 11.78 | | | 11.78 | | | 11.78 | | | 11.78 | | |
| **Sum C**[c] | 77.12 | ± | 1.42 | 83.44 | ± | 9.44 | 74.71 | ± | 12.90 | 49.92 | ± | 12.74 | 62.01 | ± | 6.01 |
| **Sum N**[c] | 20.43 | ± | 0.65 | 22.36 | ± | 2.70 | 19.97 | ± | 4.09 | 11.87 | ± | 4.00 | 15.87 | ± | 1.87 |
| $C_{Total}$[d] | 122.56 | ± | 3.90 | 134.16 | ± | 16.22 | 119.80 | ± | 24.56 | 71.25 | ± | 24.01 | 95.20 | ± | 11.21 |
| **"Missing" C**[e] | 35.08 | ± | 1.42 | 38.85 | ± | 9.53 | 36.22 | ± | 9.39 | 17.42 | ± | 11.23 | 29.97 | ± | 5.26 |

[a] C and N content was estimated based on the stoichiometry of macromolecular pools: 50% and 16% from protein; 34% and 15.5% from RNA; 36% and 16% from DNA; 74% and 6.3% from DVchlA, respectively (Geider and La Roche 2002)

[b] Estimated C in membranes assuming 14 million lipid molecules (Waldbauer et al. 2012)

[c] Sum of C and N fg cell$^{-1}$ were calculated from macromolecules pool

[d] Calculated from all N pools, $N_{total} \times 6$ (Assuming C:N ratio of 6)

[e] Estimation of C in storage molecules ($C_{Total}$ – sum C from pools)

**Calculating the expected growth of *Prochlorococcus* based on photophysiology.**

The maximal theoretical growth rate of a photosynthetic organism, assuming all fixed carbon comes from photosynthesis (no mixotrophy), can be calculated from the product of the photon flux, the DVchlA specific absorption coefficient, the photosynthetic quantum yield and the amount of chlorophyll per unit carbon. The photon flux and the amount of chlorophyll per unit carbon were measured or assessed in this study, and are 22 μmol Q m$^{-2}$ s$^{-1}$ and 360 mg DVchlA mol C$^{-1}$ (the latter assuming 3 fg DVChlA cell$^{-1}$ and 120 fg C cell$^{-1}$, see Figure 1a and Supplementary Table S1). The specific absorption coefficient (spectrally weighted) and the photosynthetic quantum yield for MIT9312 were measured by (Moore and Chisholm 1999), and are around 0.018 m$^{2}$ mg DVchlA $^{-1}$ and 0.063 mol C mol Q$^{-1}$, respectively. The product of these numbers (after unit conversion) predicts a growth rate of 0.64 day$^{-1}$, whereas cells are actually growing (during exponential stage) at just over half that rate (0.353 day$^{-1}$± 0.0239). We note that a similar discrepancy is observed also between the growth rates observed by (Moore and Chisholm 1999) (e.g. 0.78 day$^{-1}$ for MIT9312 at ~110 μmol Q m$^{-2}$ s$^{-1}$) and the rate predicted based on the photosynthetic parameters of that study, which is about 1.88 day$^{-1}$.

We note that this calculation is only a rough estimate and is sensitive to the input parameters but is likely a lower bound of the discrepancy between the theoretical and actual growth rates. First, we used the high value of C biomass (120 fg cell$^{-1}$) estimated from our protein measurements, noting that this is at the high end of measured quotas. In contrast, the value we measured for DVchlA (3-4 fg cell$^{-1}$) is within the range of previous measurements (e.g. (Chisholm et al. 1988) measured 2 fg cell$^{-1}$). Thus, if the ratio of chlorophyll to carbon increases so does the predicted growth rate. Second, this discrepancy does not take into account the reduction in growth rate during late exponential and stationary stages. Third, while the measurements of (Moore and Chisholm 1999) include also exuded C (they were not performed on filtered cells but rather on the entire culture), the acidification step required to remove the inorganic $^{14}$C may also remove some volatile forms of organic carbon, although the magnitude of this potential effect is unknown (*Prochlorococcus* have been shown to produce volatile compounds such as isoprene, but this compound is produced at an estimate of 60-100×10$^{-6}$ fg cell$^{-1}$ day$^{-1}$ and thus is a very minor C flux (Shaw et al. 2003)).

**A mathematical model of a generalized phytoplankton resolving key macromolecular pools**

The model, depicted in Supplementary Figure S3, describes the uptake of nitrogen and the uptake of carbon through photosynthesis and their conversion into various biomolecules. There are 22 parameters, listed in Supplementary Table S2, and eight variables. Five of the variables, each with the symbol $X$, relate to the amount of N per cell in the different macromolecular pools: storage ($X_{N,stor}$ in pmol N cell$^{-1}$), protein ($X_{N,prot}$ in pmol N cell$^{-1}$), DNA ($X_{N,DNA}$ in pmol N cell$^{-1}$), RNA ($X_{N,RNA}$ in pmol N cell$^{-1}$) and chlorophyll ($X_{N,Chl}$ in pmol N cell$^{-1}$). $N_{med}$ is the amount of N in the medium (dissolved inorganic N, DIN, in $\mu$M), $X_{TOC}$ represents the total organic carbon (in pmol C cell$^{-1}$), which includes both the intracellular carbon and the extracellular dissolved organic carbon (DOC), and $\rho$ is the cell density (in cells $\mu$l$^{-1}$).

**DIN uptake** from the medium follows Michaelis-Menten kinetics:

$$\frac{dN_{med}}{dt} = -V_{max,N}\rho\frac{N_{med}}{K_{Nmed}+N_{med}} \qquad (1)$$

The maximum DIN uptake rate ($V_{max,N}$) is estimated by fitting the model the data (see below for details). However, it is not possible to estimate the half-saturation constant of DIN uptake ($K_{Nmed}$) directly from the data, since that would require very precise measurements of the uptake rate as a function of the DIN concentration at the transition point from the exponential to the stationary growth stage. We obtain a value for $K_{Nmed}$ by assuming that the uptake of DIN from the medium is diffusion-limited at the lowest concentrations. Under diffusion limitation, the rate of nutrient uptake rate for a spherical cell is:

$$\frac{dN_{med}}{dt} = -4\pi r D \rho N_{med} \qquad (2)$$

with $r$ the cell radius and $D$ the diffusion coefficient of the nutrient (Berg and Purcell 1977). For $N_{med} \downarrow 0$, combining equations (1) and (2) gives $V_{max,N}\rho\frac{N_{med}}{K_{Nmed}} = 4\pi r D\rho N_{med}$, leading to $K_{Nmed} = \frac{V_{max,N}}{4\pi r D}$. Importantly, while the value of the half-saturation constant is critical for understanding nutrient limitation, models of batch culture are relatively insensitive to it, since the concentration of inorganic nutrients typically approach this value only for a short time before nutrients become depleted (Grossowicz et al. 2017).

**Dynamics of macromolecular pools:** DIN is taken up into the cellular storage, from which nitrogen is taken for the synthesis of functional macromolecules. Underlying our model is the assumption that the aggregate reactions leading to the production of these macromolecules exhibit Michaelis-Menten kinetics with storage nitrogen in the role of substrate. In addition, the amount of each cellular compound becomes distributed over a larger volume as the organisms grow and divide, a process referred to as "dilution by growth" (Kooijman 2000). We assume that proteins act as enzymes in the synthesis of RNA, DNA and Chl. Furthermore, we account for exudation of macromolecules. Putting biosynthesis, exudation, and dilution by growth together leads to the following generic equation:

$$\frac{dX_A}{dt} = \underbrace{V_{max,A}X_{N,prot}\frac{X_{N,Stor}}{K_A + X_{N,stor}}}_{Biosynthesis} - \underbrace{ex_A X_A}_{Exudation} - \underbrace{\mu X_A}_{Dilution} \qquad (3)$$

with $X_A$ a shorthand for $X_{N,RNA}$, $X_{N,DNA}$, or $X_{N,Chl}$, $V_{max,A}$ a shorthand for $V_{max,RNA}$, $V_{max,DNA}$, or $V_{max,Chl}$ (maximum synthesis rates of RNA, DNA and Chl), $K_A$ a shorthand for $K_{RNA}$, $K_{DNA}$, or $K_{Chl}$ (half-saturation constants for RNA, DNA and Chl), $ex_A$ a shorthand for $ex_{prot}$, $ex_{RNA}$, $ex_{DNA}$, or $ex_{Chl}$ (exudation rates for protein, RNA, DNA and Chl), and $\mu$ the population growth rate. For protein synthesis, we use a slightly different formulation than for the synthesis of RNA, DNA, and Chl. Most of the RNA is in the ribosomes (Bremer and Dennis, 1996) and is thus connected with protein synthesis. Indeed, an increase of the RNA:protein ratio with increasing growth rate is observed across many organisms (Scott et al. 2010). Therefore, we assume that RNA acts as an enzyme in protein synthesis:

$$\frac{dX_{N,prot}}{dt} = \underbrace{V_{max,P} X_{N,RNA} \frac{X_{N,Stor}}{K_{prot}+X_{N,stor}}}_{Biosynthesis} - \underbrace{ex_P X_{N,prot}}_{Exudation} - \underbrace{\mu X_{N,prot}}_{Dilution} \quad (4)$$

with $X_{N,prot}$ the protein per cell in N-moles, $V_{max,prot}$ the maximum protein synthesis rate per amount of RNA, and $K_P$ the half-saturation constant for protein synthesis. Our model does not account for changes in the number of genome copies per cell (e.g. (Sukenik et al. 2012) ), because the amount of DNA per cell changes relatively little throughout the experiment (Figure 1). Setting $\frac{dX_{N,DNA}}{dt}$ to 0, equation (1) gives an expression for the population growth rate:

$$\mu = \frac{V_{max,DNA}\frac{X_{N,Stor}}{K_{DNA}+X_{N,stor}}}{X_{N,DNA}} - ex_{DNA}.$$

Now that we have formulated the rates of synthesis of functional macromolecules, we can write down an equation for the storage N budget:

$$\frac{dX_{N,stor}}{dt} = \underbrace{V_{max,N} \frac{N_{med}}{K_{Nmed}+N_{med}}}_{Uptake} - \underbrace{Biosyn_{tot}}_{Biosynthesis} - \underbrace{\mu X_{N,stor}}_{Dilution} \quad (5)$$

with the total biosynthesis defined as: $Biosyn_{tot} \equiv X_{N,prot}\left(\frac{V_{max,RNA}X_{N,Stor}}{K_{RNA}+X_{N,stor}} + \frac{V_{max,DNA}X_{N,Stor}}{K_{DNA}+X_{N,stor}} + \frac{V_{max,Chl}X_{N,Stor}}{K_{Chl}+X_{N,stor}}\right) + X_{N,RNA}\frac{V_{max,prot}X_{N,stor}}{K_{prot}+X_{N,stor}}$.

The total organic carbon is determined by the balance between photosynthesis and respiration:

$$\frac{dX_{TOC}}{dt} = \underbrace{X_{N,Chl}Y(t)}_{Photosynthesis} - \underbrace{\left(r_0 X_{C,func} + r_1 Biosyn_{tot}\right)}_{Respiration} \qquad (6)$$

in which $Y(t)$ is the photosynthesis yield (in mol C (N-mol Chl)$^{-1}$ d$^{-1}$) that is allowed to vary over time. For each macromolecular pool there are two associated respiration terms, one related to maintenance of the functional macromolecules ("maintenance respiration", ($r_0 X_{C,func}$), and one related to the C cost of growth or biosynthesis ($r_1 Biosyn_{tot}$). $X_{C,func}$ is the amount of carbon in functional macromolecules (in pmol C cell$^{-1}$) and is calculated based on the elemental ratios of the various macromolecules: $X_{C,func} \equiv C{:}N_{prot}X_{N,prot} + C{:}N_{RNA}X_{N,RNA} + C{:}N_{DNA}X_{N,DNA} + C{:}N_{Chl}X_{N,Chl}$

**Parameter estimation:** We use the same Metropolis algorithm as in (Omta et al. 2017) for our parameter estimate (Supplementary Table S2). From the measurements on *Prochlorococcus* reported in this article, we estimate the maximum DIN uptake rate ($V_{mN}$), the maximum synthesis rates of protein, RNA, DNA, and Chl ($V_{max,prot}$, $V_{max,RNA}$, $V_{max,DNA}$, and $V_{max,Chl}$), the exudation rates of protein, RNA, DNA, and Chl ($ex_{prot}$, $ex_{RNA}$, $ex_{DNA}$, and $ex_{Chl}$), and the amount of DNA per cell ($X_{N,DNA}$) that is assumed constant. The half-saturation constants for the conversion of storage N into macromolecules ($K_{prot}$, $K_{RNA}$, $K_{DNA}$, $K_{Chl}$) cannot be estimated and are taken to be very small, equal to 1 fmol cell$^{-1}$. However, these parameter values have very little impact on the fits to the data. We account for uncertainty in the initial conditions by adding Gaussian white noise with a standard deviation equal to the reported measurement error to the initial concentrations.

**Model results:** Model fits for the cell density and protein, RNA, and Chlorophyll per cell are shown in Supplementary Figure S4. In each Figure panel, 100 randomly drawn simulations with parameter sets accepted by the algorithm are shown. The widths of the composite curves reflect the uncertainties in the fits, according to the algorithm. For a good fit of the total organic carbon (TOC), the photosynthetic yield (Y) needs to vary over time. Using a photosynthetic yield that is constant in time, the model does not capture the rapid increase in TOC between day 10 and day 13 of the experiment (Supplementary Figure S5a). The fit improves significantly if we allow the photosynthetic yield to vary over time (Supplementary Figure S5b). However, the model then indicates an unexplained maximum in the photosynthetic yield around day 12/13 of the experiment

(Supplementary Figure S5c). This phenomenon – an increase in photosynthetic efficiency in cells entering chlorosis (potentially caused by a reduction in shelf-shading, (Felcmanová et al. 2017)) - has not, to the best of our knowledge, been described in previously, and is a prediction of the model that can be tested in future studies.  An alternative explanation is that the respiration rates (either the maintenance respiration or the growth-related respiration) may change during batch culture, dropping as the cells become chlorotic. This will potentially result in an accumulation of DOC. In support of this possibility, the respiration rate of *Prochlorococcus* has recently been shown to change in response to incubation with dissolved organic carbon (phage lysate), without a change in growth rate (Fang et al. 2019). The current experimental data does not allow to differentiate between an increase in photosynthetic yield and a decrease in one or both of the respiration terms, and thus this question should be addressed in future studies. Additionally, the current model describes the *Prochlorococcus* community as homogenous in terms of cell physiology, and thus does not take into account the differences between chlorotic and non-chlorotic cells. Incorporating this phenomenon into mathematical models of cell physiology is an important challenge for future studies.

**Supplementary Table S2:** Model parameters

| Parameter | Description | Value | Units |
|---|---|---|---|
| $V_{mP}$ | Maximum protein synthesis rate | 9.87 | mol (N) protein/(mol (N) RNA $d^{-1}$) |
| $V_{mR}$ | Maximum RNA synthesis rate | 0.12 | mol (N) RNA/(mol (N) protein $d^{-1}$) |
| $V_{mD}$ | Maximum DNA synthesis rate | 0.006 | mol (N) DNA/(mol (N) protein $d^{-1}$) |
| $V_{mChl}$ | Maximum Chl synthesis rate | 0.028 | N-mol Chl/(N-mol protein $d^{-1}$) |
| $V_{mN}$ | Maximum DIN uptake rate | 2.38 | $\mu$M $d^{-1}$ |

| Parameter | Description | Value | Units |
|---|---|---|---|
| $X_{N,DNA}$ | Cellular DNA content | 6.49 | amol (N) cell$^{-1}$ |
| $ex_P$ | Protein exudation | 0.14 | d$^{-1}$ |
| $ex_R$ | RNA exudation | 0.65 | d$^{-1}$ |
| $ex_D$ | DNA exudation | 0.14 | d$^{-1}$ |
| $ex_{Chl}$ | Chl exudation | 1.07 | d$^{-1}$ |
| $K_P$ | Half-saturation constant for protein synthesis | 1.0 | fmol (N) cell$^{-1}$ |
| $K_R$ | Half-saturation constant for RNA synthesis | 1.0 | fmol (N) cell$^{-1}$ |
| $K_D$ | Half-saturation constant for DNA synthesis | 1.0 | fmol (N) cell$^{-1}$ |
| $K_{Chl}$ | Half-saturation constant for Chl synthesis | 1.0 | fmol (N) cell$^{-1}$ |
| $r_0$ | Maintenance respiration rate | 0.1 | d$^{-1}$ |
| $r_1$ | Growth respiration | 2.0 | mol C/(mol N) |
| $C:N_{Pro}$ | C:N ratio of protein | 4.0 | mol C/(mol N) |
| $C:N_{RNA}$ | C:N ratio of RNA | 3.0 | mol C/(mol N) |
| $C:N_{DNA}$ | C:N ratio of DNA | 3.0 | mol C/(mol N) |
| $C:N_{Chl}$ | C:N ratio of Chl | 13.0 | mol C/(mol N) |
| $D$ | Diffusion coefficient of ammonia | $2 \times 10^{-9}$ | m$^2$/t |

| Parameter | Description | Value | Units |
|---|---|---|---|
| $r$ | Cell radius | 0.8 | $\mu m$ |

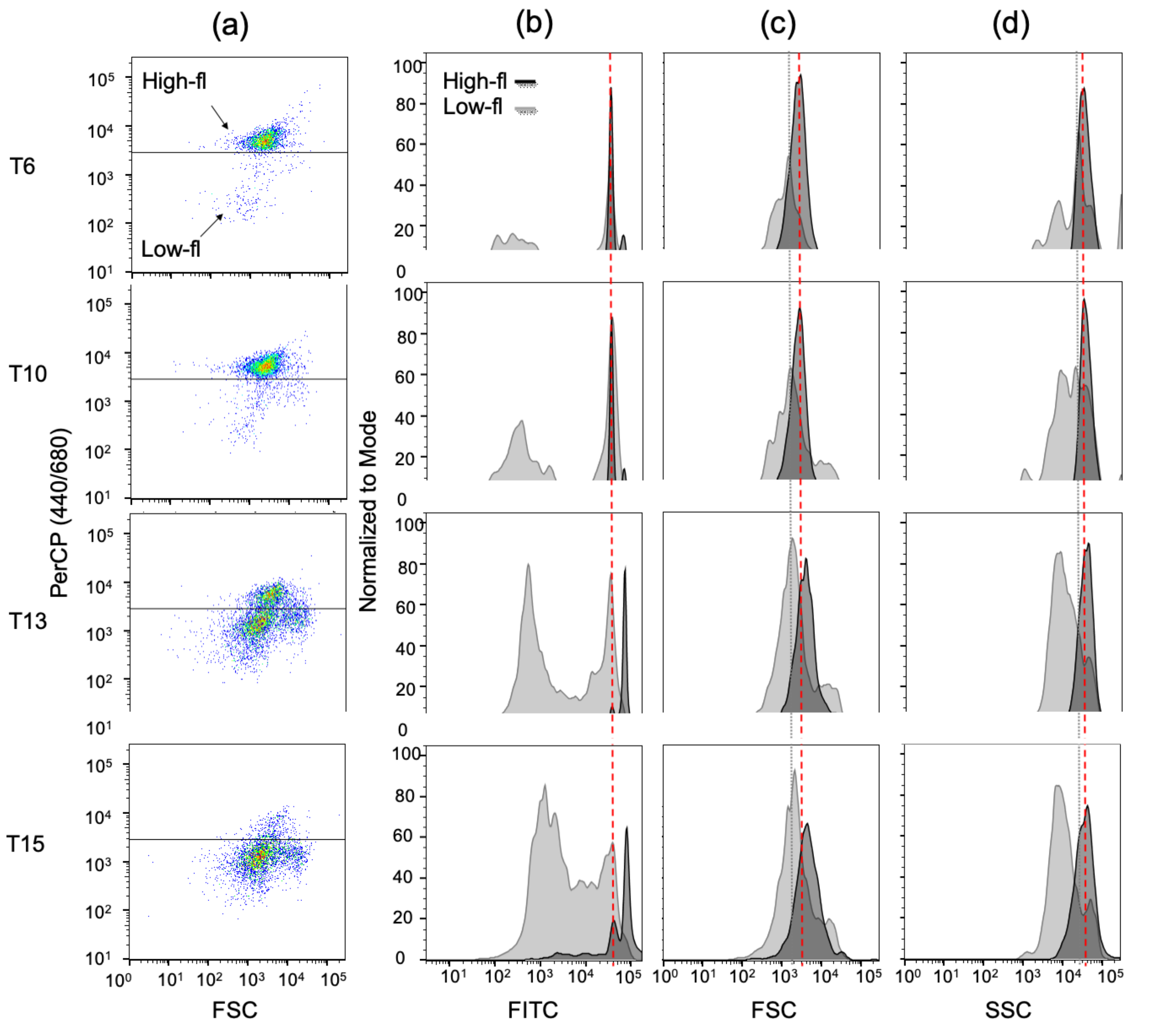

**Supplementary Figure S1: Changes to flow-cytometrically defined parameters in chlorotic and non-chlorotic cells during batch culture.** a) Scattergrams of forward scatter (FSC, a proxy for cell size) vs chlorophyll autofluorescence, showing the emergence of low-fl populations (PerCP 440/680). b) low-fl cells are more weakly stained by Sybr Green, suggesting a decrease in the cell content of nucleic acids. In addition, the high-fl population exhibited a bimodal Sybr-green staining, with the higher staining potentially representing cells with two chromosomes (G2). During late stationary and decline stages, most of the cells in the high-fl population had higher Sybr staining, which we speculate might be due to cell cycle arrest following N starvation. c) Forward Scatter (FSC) of both high-fl and low-fl cells increases during late stationary and decline stages, consistent with cell cycle arrest and/or the accumulation of stored carbon. d) low-fl cells have significantly lower side scatter (SSC). In eukaryotic cells SSC is related to the granularity or optical complexity of the cells, although this is less clear in small cells such as *Prochlorococcus*, whose size is close to the wavelength of the laser light. Nevertheless, we speculate that these changes may be due to the degradation of the photosynthetic pigments or the thylakoid membranes. The dashed and dotted lines in panels b-d show the mode of the high-fl and low-fl cells at t=0, enabling a direct visual comparison between the time points.

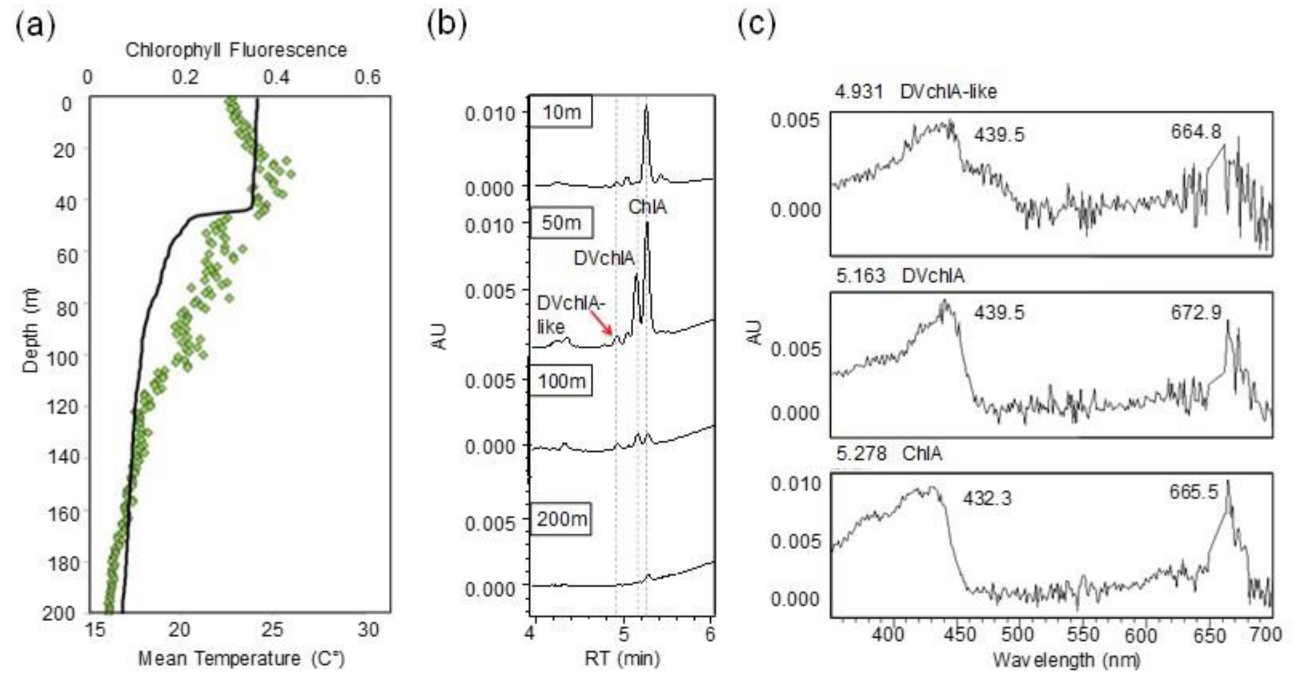


**Supplementary Figure S2: Observation of a DVchlA-like pigment in the Eastern Mediterranean Sea.** a) Oceanographic profiles of temperature and chlorophyll fluorescence (binned at one-meter intervals) from station n-1200 in the Eastern Mediterranean Sea during November 2015 (Manuscript in prep). b) Sections of UPLC chromatograms of samples collected at different depths, showing an increase in the amount of DVchlA with depth (consistent with an increase in *Prochlorococcus* cell numbers observed by flow cytometry, now shown). A DVchlA-like pigment with a similar retention time to that shown in Figure 2c is highlighted. c) Absorbance spectra of DVchlA, the DVChlA-like pigment and ChlA from the chromatograms at a depth of 50m shown in panel b.

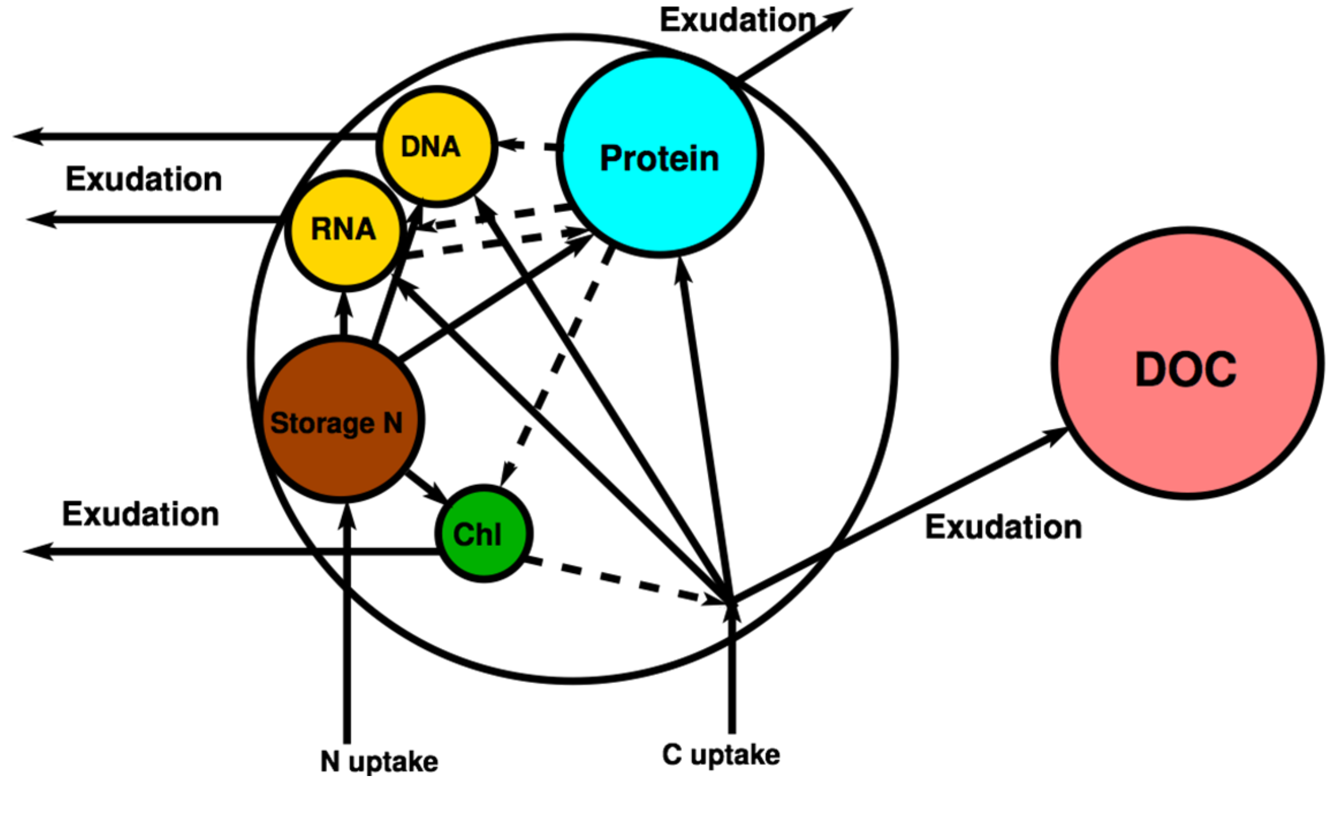


**Supplementary Figure S3: A schematic depiction of the model cells with the different compartments, each representing a major macromolecular pool**. Solid lines indicate material fluxes, dashed lines indicate enzymatic action.

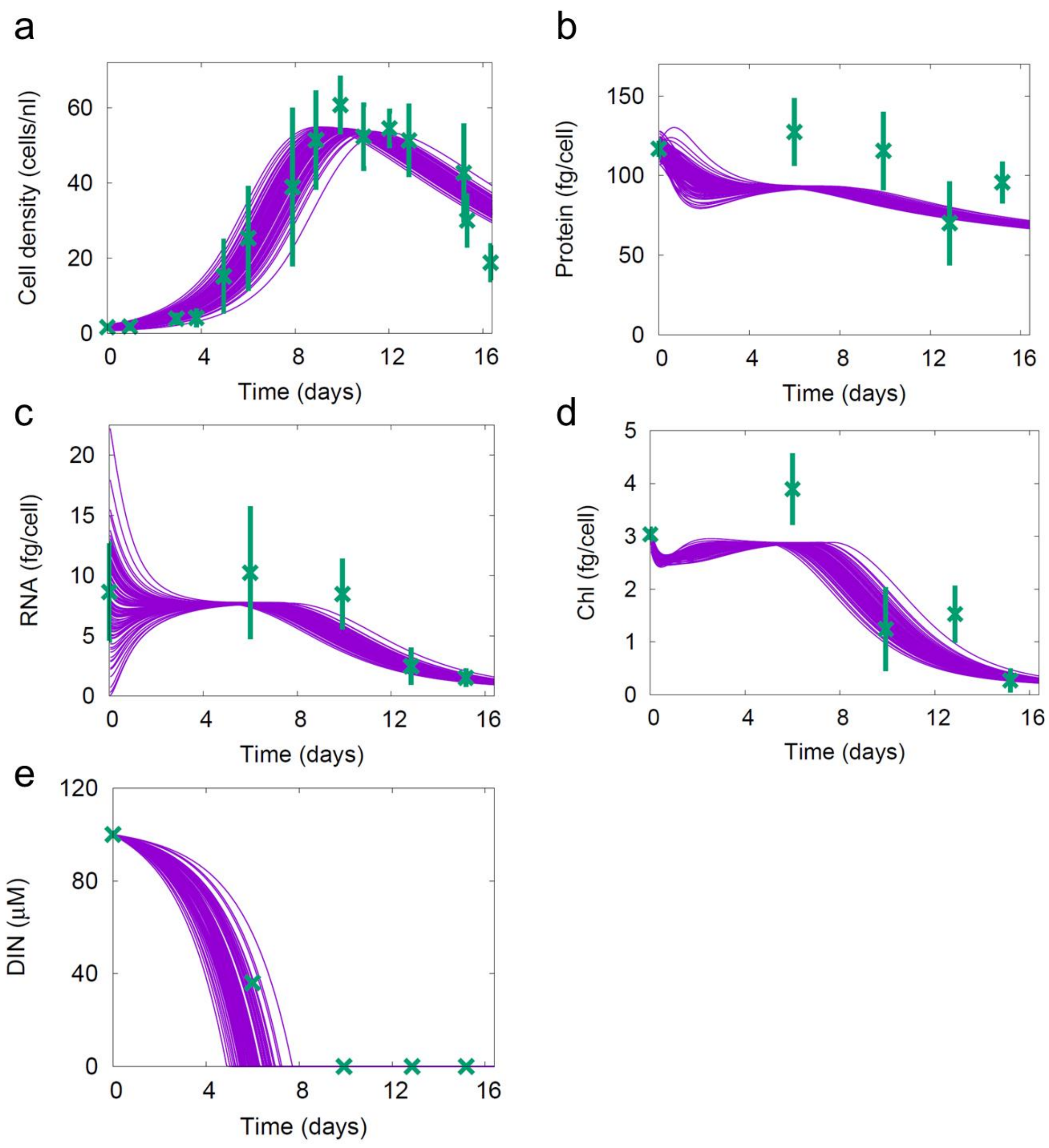


**Supplementary Figure S4: Model fits for cell density (a), protein per cell (b), RNA per cell (c), DVchlA per cell (d) and inorganic N ($NH_4$) in the media (e).** Purple lines are 100 randomly selected model runs, and are compared to experimental results (green, same as in Figure 1).

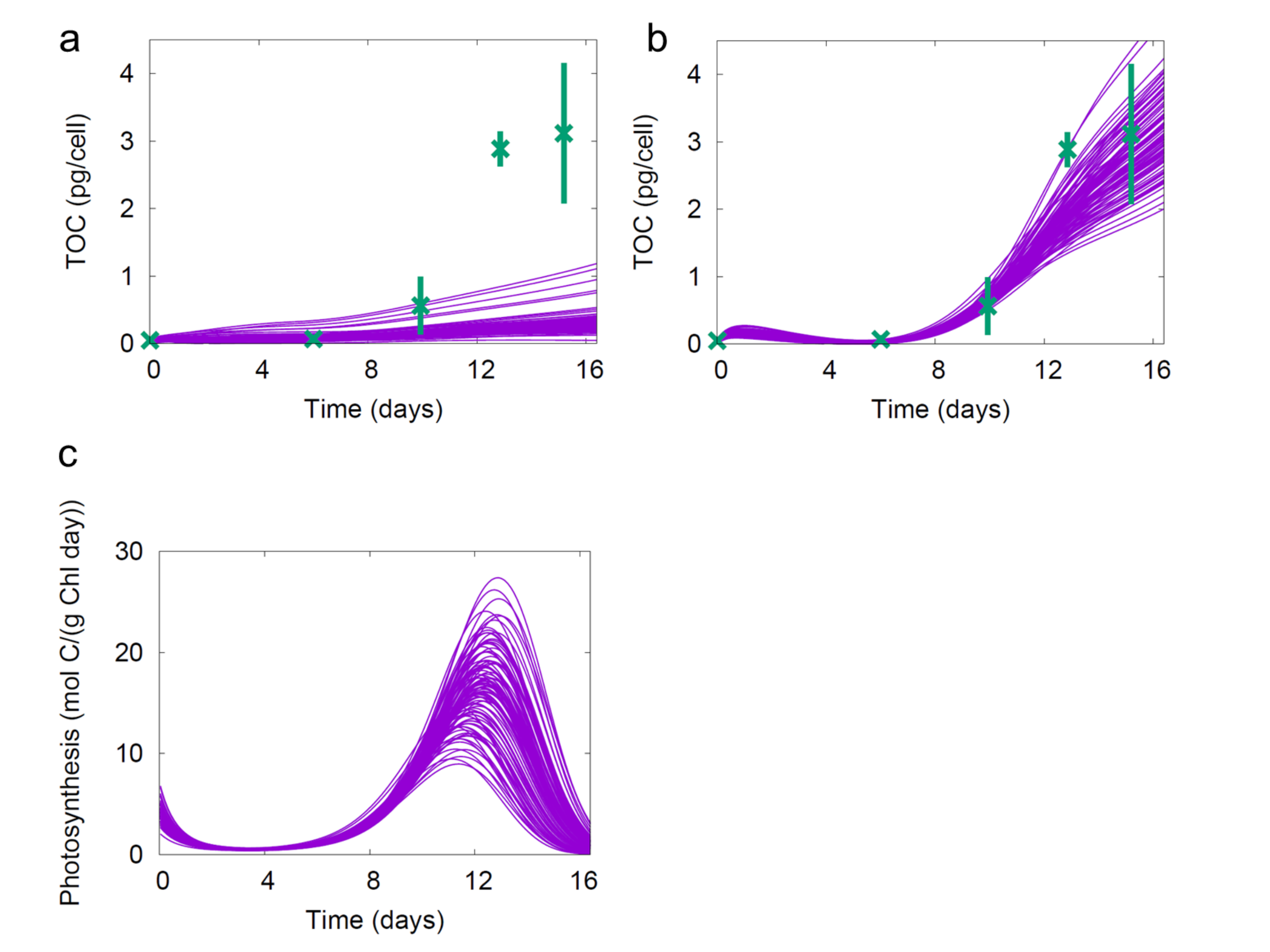


**Supplementary Figure S5: Photosynthetic yield needs to increase in chlorotic cells for the model to reproduce high DOC production.** Panels a and b show the modeled TOC (which includes both the cellular C and DOC) modelled when photosynthesis is constant (a) and when it is allowed to vary (b). Panel c shows the photosynthesis rates required for the modelled TOC curves shown in panel b. Purple lines are 100 randomly selected model runs, and are compared to experimental results (green, same as in Figure 1).
